# Cross-kingdom conservation of Arabidopsis RPS24 function in 18S rRNA maturation

**DOI:** 10.1101/2023.04.21.537868

**Authors:** Adrián Cabezas-Fuster, Rosa Micol-Ponce, Raquel Sarmiento-Mañús, María Rosa Ponce

## Abstract

All 81 ribosomal proteins (RPs) that form the Arabidopsis (*Arabidopsis thaliana*) 80S ribosome are encoded by several paralogous genes. For example, the nearly identical RPS24A and RPS24B proteins are encoded by *RPS24A* and *RPS24B*, respectively. Here we explored the functions of RPS24A and RPS24B in Arabidopsis. Their encoding genes exhibit combined haploinsufficiency, as at least two wild-type copies of either *RPS24A* or *RPS24B* are required for plant viability and at least three are required for normal plant development. Loss-of-function of either gene caused a pointed-leaf phenotype, a typical phenotype of null or hypomorphic recessive alleles of genes encoding ribosome biogenesis factors (RBFs) or RPs. We also found that RPS24A and RPS24B act as RBFs during early stages of 18S ribosomal RNA (rRNA) maturation, as loss of RPS24A or RPS24B function reduced the 18S/25S rRNA ratio. An RPS24B-GFP fusion protein predominantly localized to the nucleolus, as expected. The *rps24b-2* mutation strengthened the phenotypes of the RBF mutants *mRNA transporter4-2* and *small organ4-3*, which are defective in 5.8S rRNA maturation. This synergistic interaction might be an effect of increased 45S rDNA transcription, which we also observed in the *rps24* mutants. Therefore, the Arabidopsis RPS24 proteins act as RBFs during 18S rRNA maturation, like their human and yeast putative orthologs. Only two plant RPs were previously shown to act not only as structural components of the ribosome but also as RBFs. We provide evidence that RPS24 proteins also regulate 45S rDNA transcription, which has not been described for their yeast or human orthologs.

## INTRODUCTION

The 80S cytoplasmic ribosome (hereafter, the ribosome) of Arabidopsis (*Arabidopsis thaliana*) is composed of four ribosomal RNAs (rRNAs) and 81 ribosomal proteins (RPs; Wilson and Cate, 2012), consisting of 33 RPs in its small (40S) subunit and 48 in its large (60S) subunit (Barakat et al., 2001). In plants, rRNAs are encoded by 5S and 45S rDNA genes, which are transcribed in the nucleoplasm and the nucleolus by RNA polymerase III or RNA polymerase I, respectively. All eukaryotic genomes include hundreds of rDNA genes arranged in tandem in several loci. Each 45S rDNA gene is composed (from its 5′ to 3′ end) of a promoter and a transcriptional unit that contains the 5′ external transcribed spacer (5′-ETS), the 18S, 5.8S and 25S rRNA sequences (in plants and yeast; 28S in animals), and a 3′-ETS. These three regions encoding rRNAs are separated by two internal transcribed spacers: ITS1 is located between the sequences of the 18S and 5.8S rRNAs, while ITS2 is located between the sequences of the 5.8S and 28S/25S rRNAs (Supplemental Figure 1).

The processing of 45S (47S in animals and 35S in yeast [*Saccharomyces cerevisiae*]) pre-rRNA is a multistep procedure involving more than 100 ribosome biogenesis factors (RBFs), which carry out endo- and exo-nucleolytic cleavages and chemical modifications to generate 28S/25S, 18S and 5.8S mature rRNAs. This intricate rRNA production consists of two partially redundant pathways, which are named according to the region of the early 35S(P) pre-rRNA in which the first endonucleolytic cleavage occurs: the 5′-ETS-first and ITS1-first pathways. There is an additional pathway in plants, the ITS2-first pathway (Supplemental Figure 1; Palm et al., 2019). The functions of many RBFs are partially or fully conserved across animals, fungi and plants (reviewed in Sáez-Vásquez and Delseny, 2019).

Yeast mRNA transport4 (Mtr4) is an ATP-dependent RNA helicase that acts as a cofactor of the nucleolar exosome, which has 3′→5′ exonuclease activity. Loss-of-function mutations of the yeast *Mtr4* gene impair 5.8S and 25S rRNA maturation through the ITS1-first pathway and leads to an imbalance in the ratio of 40S/60S ribosomal subunits (Thoms et al., 2015). Arabidopsis MTR4 is required for 18S and 5.8S rRNA maturation and to eliminate 5′-ETS processing by-products produced in the ITS1-first pathway (Lange et al., 2011). Yeast Mtr4 interacts with Nucleolar protein 53 (Nop53), which has orthologs in humans (Glioma Tumor-Suppressor Candidate Region 2; GLTSCR2) and Arabidopsis (SMALL ORGAN 4; SMO4), all of which are involved in equivalent steps of 5.8S rRNA maturation (Tafforeau et al., 2013; Micol-Ponce et al., 2020). Ribosomal RNA processing protein 7 (Rrp7) is another yeast RBF involved in 18S rRNA processing that functions in the ITS1-first pathway, as do its human (RRP7A) and Arabidopsis (RRP7) orthologs (Micol-Ponce et al., 2018; Farooq et al., 2020; Supplemental Figure 1).

Most RPs that form part of the 40S ribosomal subunit also function as RBFs in the nuclear and cytoplasmic steps of 18S rRNA maturation in yeast, including Ribosomal protein S24 (Rps24; Ferreira-Cerca et al., 2005). Human RPS24 and yeast Rps24 play analogous roles in 5′-ETS processing. Loss-of-function mutations of the yeast *Rps24* and human *RPS24* orthologs lead to the accumulation of 23S and 30S pre-rRNAs, respectively (which are equivalent to Arabidopsis P-A_3_ pre-rRNA) and a reduction in the amounts of human 21S and 18S-E, and yeast 21S and 20S pre-rRNAs, which are equivalent to Arabidopsis 18S-A_3_ and 20S, respectively. All of these pre-rRNAs are precursors of the 18S rRNA produced by the ITS1-first pathway (Supplemental Figure 1). Since this pathway is the major contributor to 18S rRNA production, its level is reduced by human and yeast *rps24* loss-of-function mutations (Ferreira-Cerca et al., 2005; Choesmel et al., 2008).

Arabidopsis *MORPHOLOGY OF ARGONAUTE1-52 SUPPRESSED 2* (*MAS2*) is the ortholog of the human gene encoding NF-κ-B-activating protein (NKAP), which functions in transcriptional repression and splicing in animals (Pajerowski et al., 2009; Burgute et al., 2014; Sánchez-García et al., 2015). We previously identified physical interactors of MAS2 in a yeast two-hybrid assay (Sánchez-García et al., 2015), including RRP7 (Micol-Ponce et al., 2018), SMO4 (Micol-Ponce et al., 2020) and RPS24B. Here, we studied the functions of the Arabidopsis co-orthologs of yeast *Rps24* and human *RPS24*: AT3G04920 and AT5G28060, which encode RPS24A and RPS24B, respectively. We established that both RPS24A and RPS24B act as RBFs in the ITS1-first pathway, like their human and yeast orthologs. Our results also suggest that RPS24A and RPS24B participate in the transcriptional repression of 45S rDNA, a role that has not been proposed for any of their orthologs.

## RESULTS

### Arabidopsis *RPS24A* and *RPS24B* show combined haploinsufficiency

The Arabidopsis *RPS24A* (AT3G04920) and *RPS24B* (AT5G28060) genes are very similar in structure, with 5 exons, and their protein products sharing 96% sequence identity (Supplemental Figure 2). We initially focused our study on RPS24B, since we previously showed that it interacted with MAS2 in a Y2H assay (Sánchez-García et al., 2015). Two putatively null alleles of *RPS24B*, *rps24b-1* and *rps24b-2*, were previously identified as mutants with impaired leaf development; *rps24b-1* harbors a 72-bp deletion in the coding region of its fifth exon (Horiguchi et al., 2006; Horiguchi et al., 2011), and *rps24b-2* carries a T-DNA insertion in its third intron (Figure 1A; Horiguchi et al., 2011; Wang et al., 2018). Both mutants exhibited an almost identical mild pointed-leaf phenotype (Figure 1F; Horiguchi et al., 2011; Wang et al., 2018).

**Figure 1.**
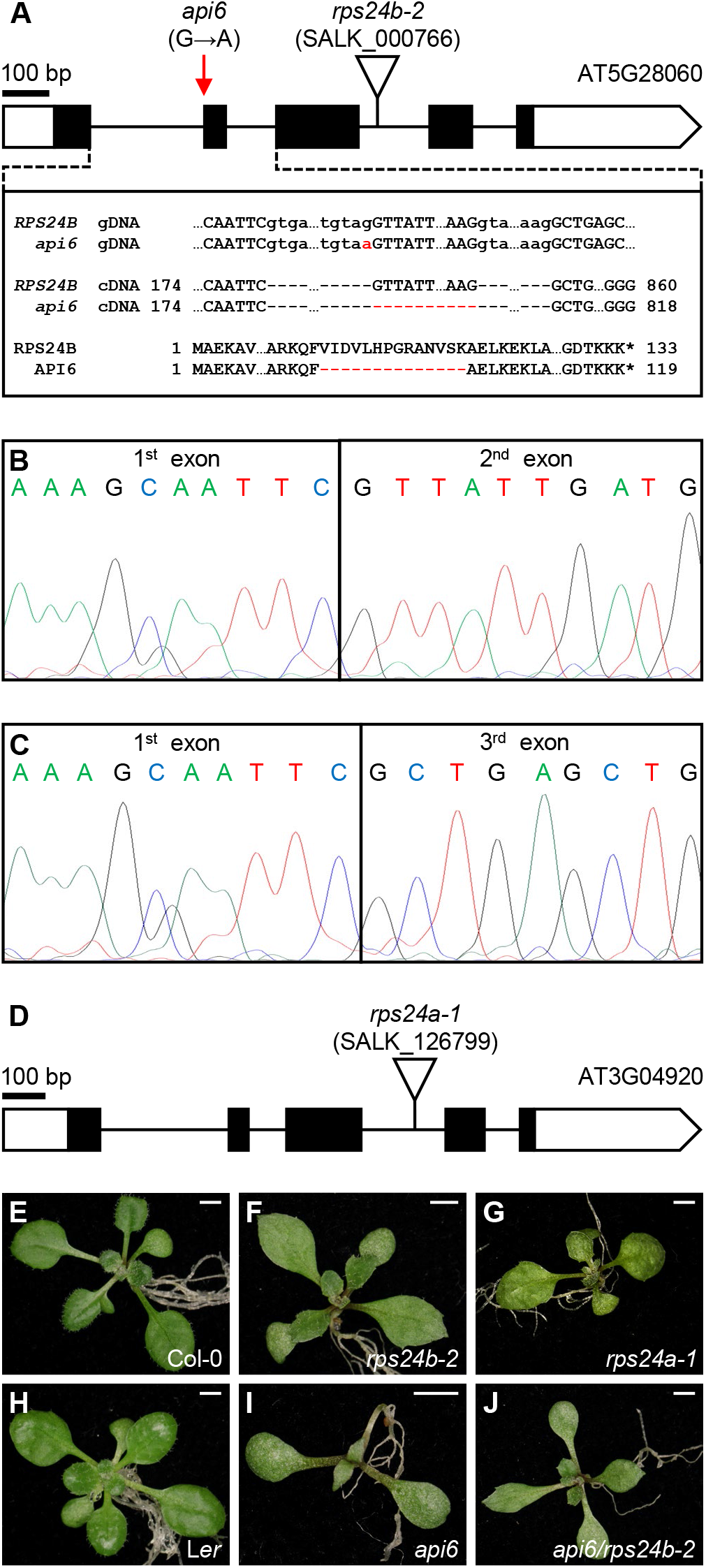
Alleles of the *RPS24A* and *RPS24B* genes studied in this work. (A) Schematic representation of the *RPS24B* locus, with the positions and nature of its mutations indicated. Exons and introns are depicted as boxes and lines, respectively, and white boxes represent untranslated regions (UTRs). A red arrow and a triangle mark the point mutation of *api6* and the T-DNA insertion of *rps24b-2*, respectively. The molecular changes caused by *api6* and *rps24b-2* are shown in red in the corresponding cDNAs and predicted proteins. (B and C) Electropherograms showing the cDNA sequences of *RPS24B* in L*er* (B) and *api6* (C). Total RNA was extracted from seedlings collected 15 days after stratification (das). (D) Structure of the *RPS24A* gene, represented as described in A. (E-J) Rosettes of Col-0 (E), *rps24b-2* (F), *rps24a-1* (G), L*er* (H), *api6* (I) and *api6/rps24b-2* F_1_ seedlings (J). Scale bars, 2 mm. Photographs were taken 14 das.

The *apiculata6* (*api6*) mutant was previously isolated in a large-scale screen for leaf morphological mutants from an EMS mutagenized population in the L*er* background (Berná et al., 1999). Using iterative linkage analysis (Ponce et al., 1999; Ponce et al., 2006), we mapped the *api6* mutation to a genomic region containing 49 genes flanked by the molecular markers cer449133 (between AT5G27905 and AT5G27910) and cer451402 (in AT5G28200; Supplemental Table 1). Sanger sequencing of AT5G28060 (*RPS24B*), the most plausible candidate gene, revealed an EMS-type G→A transition in the last nucleotide of the first intron (Figure 1A). Since this nucleotide forms part of a 3′ splicing site (3′SS) of the first intron, the *api6* mutation is predicted to cause the absence of the second exon of the gene—which is 42-nt in length—from the mature mRNA of *api6* (Figure 1B and C). The protein produced by the *api6* mutant is predicted to lack 14 residues present in wild-type RPS24B (amino acids 25-38; Supplemental Figure 2). An *api6 × rps24b-2* cross confirmed that these mutants are allelic (Figure 1E, F, H-J). Like many mutants in the L*er* genetic background, the leaf phenotype of *api6* was stronger than that of *rps24b-1* or *rps24b-2*, which are in the Col-0 background (Pérez-Pérez et al., 2009).

We also studied SALK_126799, a line that carries a T-DNA insertion in the third intron of *RPS24A,* which we named *rps24a-1* (Figure 1D and G). Homozygous *rps24a-1* plants were viable and exhibited a pointed-leaf phenotype, which was milder than that of *rps24b-2* and *api6* (Figure 1E-I). RPS24A and RPS24B exhibited similar protein abundance patterns, as determined in the Arabidopsis THaliana ExpressioN Atlas (Athena) database (http://athena.proteomics.wzw.tum.de:5002/master_arabidopsisshiny/; Mergner et al., 2020), with similar levels throughout all organs and developmental stages, with a few exceptions (Supplemental Dataset 1 [highlighted in red]). The almost constitutive expression of *RPS24A* and *RPS24B* (at a lower level compared to other genes encoding RPs) was also described by Savada and Bonham-Smith (2014) using microarray data from the GENEVESTIGATOR platform (Hruz et al., 2008).

Since the mutant phenotypes of *rps24a-1* and *rps24b-2* pointed to the functional redundancy of *RPS24A* and *RPS24B*, we performed *rps24a-1 × rps24b-2* reciprocal crosses. All *RPS24A/rps24a-1;RPS24B/rps24b-2* F_1_ plants exhibited a mutant phenotype indistinguishable from that of their parents, suggesting non-allelic non-complementation (Figure 2A-D). Selfing of such diheterozygotes generated F_2_ progeny with only two phenotypic classes: 221 wild-type and 161 mutant (indistinguishable from their parents) plants. We genotyped 59 F_2_ phenotypically mutant plants, finding 10 *rps24a-1/rps24a-1;RPS24B/RPS24B*, 17 *RPS24A/RPS24A;rps24b-2/rps24b-2* and 32 *RPS24A/rps24a-1;RPS24B/rps24b-2* plants. We identified no *rps24a-1/rps24a-1;RPS24B/rps24b-2* or *RPS24A/rps24a-1;rps24b-2/rps24b-2* sesquimutants or double mutants among the F_2_ progeny (Figure 2A-D); the absence of these genotypes suggests their lethality. In agreement with the proposed early lethality of the genotypes with fewer than two wild-type doses of either *RPS24A* or *RPS24B* (expected to be 31.25% of seeds), the siliques of several diheterozygous F_1_ plants exhibited a rate of 27% aborted or unfertilized ovules compared to Col-0 (Figure 2E-H).

**Figure 2.**
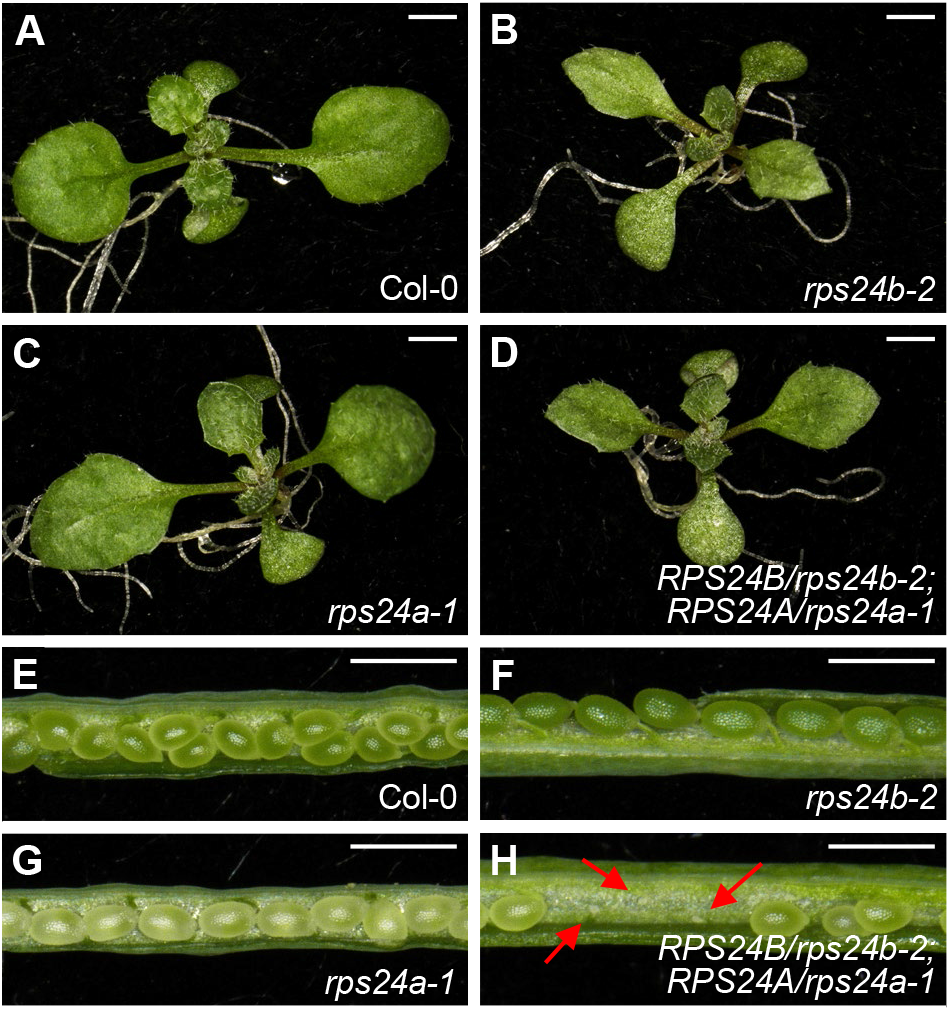
Genetic evidence for the combined haploinsufficiency of *RPS24A* and *RPS24B*. (A-D) Rosettes of Col-0 (A), *rps24b-2* (B), *rps24a-1* (C) and (D) *RPS24B/rps24b-2;RPS24A/rps24a-1* plants. (E-H) Siliques of (E) Col-0, (F) *rps24b-2*, (G) *rps24a-1* and (H) *RPS24B/rps24b-2;RPS24A/rps24a-1* plants. Scale bars, 2 mm (A-D), or 1 mm (E-H). Photographs were taken (A-D) 14 or 39 (E-H) das.

To determine whether *rps24a-1*, *rps24b-2* and *api6* were hypomorphic or null alleles and to test the existence of any dosage compensation mechanism for the expression of *RPS24A* or *RPS24B*, we analyzed *RPS24B* mRNA levels by RT-PCR. We used primers upstream and downstream of the T-DNA insertion of *rps24b-2* (Supplemental Figure 3 and Supplemental Table 2). *RPS24B* mRNA levels were very similar in *api6* and L*er*, but *RPS24B* mRNA levels were undetectable in *rps24b-2* and were not higher in *rps24a-1* than in the wild type (Supplemental Figure 3). We did not determine whether *rps24a-1* was null, because it was impossible to design RT-PCR primers specific for this allele that would amplify the region flanking its T-DNA insertion, due to its sequence similarity with *rps24b-2*. However, the phenotype of the single mutants and the lethality of the reciprocal sesquimutants strongly suggest that *rps24a-1* and *rps24b-2* are null alleles.

### RPS24B predominantly localizes to the nucleolus

Since ribosomal subunits are synthetized and assembled in the nucleolus and the nucleoplasm, but their final maturation occurs in the cytoplasm, not few RPs have been detected in all three subcellular compartments (Pendle et al., 2005; Palm et al., 2016; Montacié et al., 2017; Ayash et al., 2021). As expected from their known roles as structural components of the ribosome, Arabidopsis RPS24A and RPS24B have been found in the cytoplasm in several proteomic studies (Chang et al., 2005; Giavalisco et al., 2005; Carroll et al., 2008; Hummel et al., 2015), and their abundances appear to be similar, at least in ribosomes purified from cell cultures (Salih et al., 2020). RPS24A and RPS24B have also been localized to the nucleolus and nucleoplasm (Pendle et al., 2005; Palm et al., 2016; Montacié et al., 2017; Ayash et al., 2021).

To visualize the subcellular localization of RPS24B, we generated the *35S_pro_:RPS24B:GFP* construct and used it to transform Col-0 plants. We primarily detected green fluorescent protein (GFP) fluorescence in the nucleolus, but also in the cytoplasm (Figure 3), as observed for other duplicated Arabidopsis RPs of the small and large subunits: RPS3aA/B, RPS8A/B, RPL7aA/B, RPL15A/B and RPL23aA/B (Savada and Bonham-Smith, 2014). Three protein-localization predictor tools provided in silico support to our experimental findings with the *35S_pro_:RPS24B:GFP* transgenic plants. LOCALIZER (https://localizer.csiro.au/; Sperschneider et al., 2017) identified a putative nuclear localization signal (NLS) in the C-terminal regions of RPS24A and RPS24B, which matches the nucleolar localization signal (NoLS) found using Nucleolar localization sequence Detector (NoD) software (Supplemental Figure 2; http://www.compbio.dundee.ac.uk/www-nod/index.jsp; Scott et al., 2011). MULocDeep (https://www.mu-loc.org; Jiang et al., 2021), which is able to predict the localization of any protein in 44 suborganellar compartments, clearly predicted a nucleolar localization for both RPS24 proteins, in addition to a lower-likelihood prediction for a cytoplasmic and mitochondrial localization (Supplemental Figures 4 and 5). The predicted nucleolar localization of RPS24A and experimental confirmation for RPS24B suggest that both proteins function in early steps of ribosome biogenesis. However, although the subcellular localization of the RPS24B-GFP fusion protein was the same in transgenic plants in the Col-0 or *rps24b-*2 background, the *35S_pro_:RPS24B:GFP* transgene did not rescue the morphological defects caused by the *rps24b-2* mutation.

**Figure 3.**
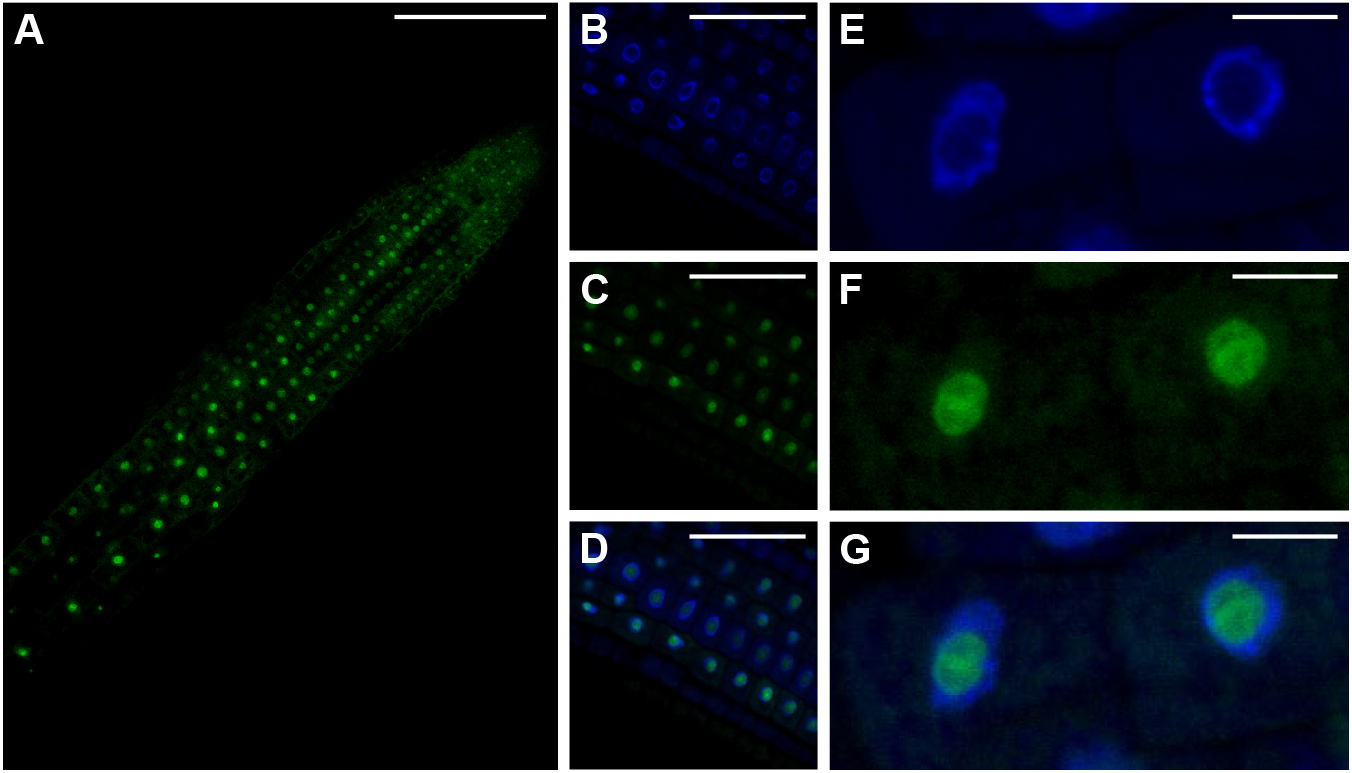
Subcellular localization of RPS24B. (A-G) Confocal laser-scanning micrographs of root cells from transgenic *35S_pro_:RPS24B:GFP* seedlings in the Col-0 background, collected 5 das. Fluorescent signals correspond to GFP (A, C, F), 4′,6-diamidino-2-phenylindole (DAPI) (B, E), and their overlay (D, G). Scale bars, 100 µm (A), 50 µm (B-D), or 10 µm (E-G).

### 45S pre-rRNA processing is delayed in the *rps24* mutants

As previously mentioned, human and yeast RPS24 act as RBFs in 18S rRNA maturation. In both species, depletion of RPS24 results in inhibited 5′-ETS processing, the accumulation of pre-rRNAs that include 5′-ETS (human 30S and yeast 23S pre-rRNAs) and reduced levels of 21S and 18S-E pre-rRNAs in humans and their corresponding 21S and 20S pre-RNAs in yeast (Supplemental Figure 1), all of which are 18S rRNA precursors (Ferreira-Cerca et al., 2005; Choesmel et al., 2008; Robledo et al., 2008).

Taken together, the current identification of RPS24B as a predominantly nucleolar factor and the previous finding of RPS24A and RPS24B in the Arabidopsis nucleolar proteome via large-scale analysis (Pendle et al., 2005; Palm et al., 2016) point to a conserved role for these proteins as RBFs. To test this hypothesis, we analyzed 45S pre-rRNA processing in the *rps24* mutants. We carried out gel blot analysis of total RNA extracted from *rps24a-1*, *rps24b-2* and *api6*, which was hybridized with the S2, S7 and S9 probes. These probes are complementary to segments of the 5′-ETS, ITS1, and ITS2 of 45S pre-rRNA, respectively, allowing several 25S, 18S and 5.8S rRNA precursors to be detected (Figure 4A). We used the *smo4-3* and *mtr4-2* mutants as controls (Lange et al., 2011; Micol-Ponce et al., 2020). Loss of SMO4 function causes nucleolar hypertrophy and the accumulation of P-A_3_, a precursor of 18S rRNA, and perturbs 5.8S rRNA maturation (Micol-Ponce et al., 2020). MTR4 is an exosome cofactor involved in the processing of 5.8S pre-rRNA (like SMO4) but also in the degradation of P-P′, a by-product derived from 5′-ETS processing (Lange et al., 2011).

**Figure 4.**
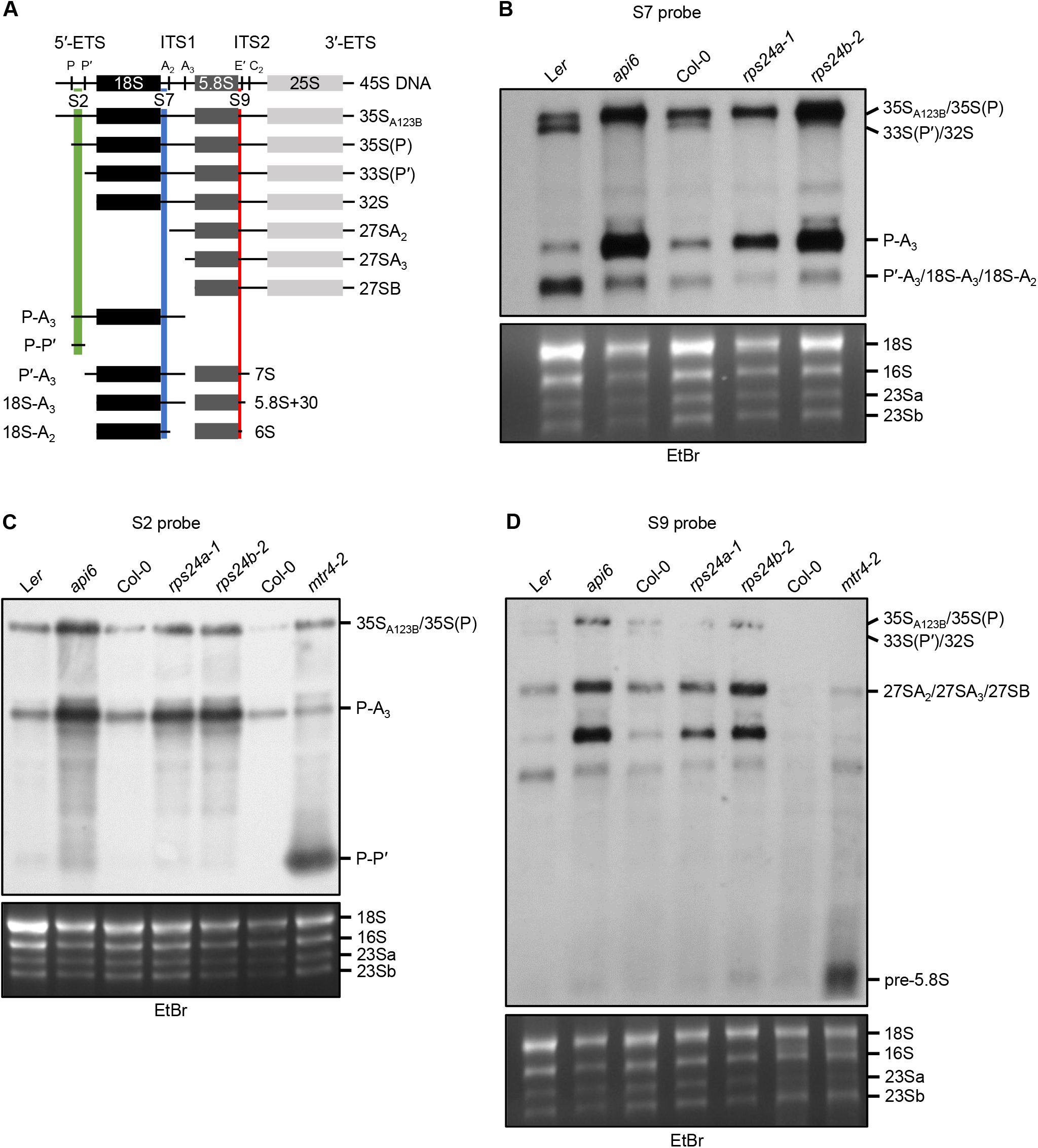
rRNA maturation in the *rps24a* and *rps24b* mutants. (A) Diagram of 45S pre-rRNA processing intermediates detected in RNA gel blots. The regions hybridizing to each probe are highlighted in green (S2 probe), yellow (S7 probe) or blue (S9 probe). Vertical bars in 45S rRNA indicate the endonucleolytic cleavage sites in pre-rRNAs that are relevant to this analysis. ETS and ITS indicate external and internal transcribed spacers, respectively. (B-D) Visualization of the processing of the 5.8S, 18S and 25S rRNA precursors by gel blot analysis. Total RNA from seedlings collected 15 das was separated on a formaldehyde-agarose gel, transferred to a nylon membrane and hybridized with the S7 (B), S2 (C), or S9 probe (D). EtBr, ethidium bromide-stained gels, visualized before blotting, serving as loading controls. Similar results were obtained in at least two independent experiments.

Using the S7 probe, we detected the accumulation of P-A_3_ and 35S_A123B_/35S(P) pre-rRNAs in *rps24a-1*, *rps24b-2* and (especially) *api6* plants compared to their respective wild types. We also detected the depletion of 33S(P′)/32S pre-rRNA and to a lesser extent P′-A_3_/18S-A_3_/18S-A_2_ species, which result from the processing of 35S_A123B_/35S(P) and P-A_3_ precursors, respectively (Figure 4B). P-A_3_ pre-rRNA is generated by the cleavage of the A_3_ site in 35S(P) pre-rRNA, the first 18S rRNA precursor in the ITS1-first pathway [Figure 4A and Supplemental Figure 1; reviewed in Sáez-Vásquez and Delseny (2019)]. We confirmed the accumulation of 35S_A123B_/35S(P) and P-A_3_ pre-rRNAs using the S2 probe, but we only observed the accumulation of the P-P′ species in *mtr4-2* (Figure 4C), as previously described (Lange et al., 2011). Using the S9 probe, we did not find any alteration in 5.8S rRNA maturation in any of the three mutants studied (Figure 4D).

Reduced levels of the 21S and 18S-E precursors of 18S pre-rRNA (Supplemental Figure 1) associated with the lack of RPS24 activity in human cells result in a 40% reduction in mature 18S rRNA levels (Choesmel et al., 2008; Robledo et al., 2008). Similar effects were found in the absence of RPS24 function in yeast: 35S, 32S and 23S pre-rRNAs accumulated, 20S pre-rRNA could not be detected, and the levels of mature 18S rRNA were reduced by 95% compared to the wild type (Ferreira-Cerca et al., 2005). The impaired 18S pre-rRNA processing due to the loss of RPS24A and RPS24B function in Arabidopsis suggests that the levels of mature 18S rRNA might be reduced in these mutants. Therefore, we analyzed the rRNA profiles of the mutants using a bioanalyzer. The 18S/25S rRNA ratio was reduced by 4% in the *rps24a-1* mutant, by 27% in *rps24b-2* and by 32% in *api6* compared to their corresponding wild types (Supplemental Figure 6). These results provide further evidence for a conserved role for RPS24A and RPS24B in the early steps of 18S rRNA maturation, particularly in the cleavage of sites internal to the 5′-ETS, as already observed for their yeast and human putative orthologs (Ferreira-Cerca et al., 2005; Choesmel et al., 2008).

### The *rps24b-2* mutation enhances the morphological and molecular defects of mutants in genes encoding RBFs that function in 18S and 5.8S rRNA maturation

To genetically confirm the involvement of RPS24 in ribosome biogenesis, we obtained double mutant combinations between *rps24b-2* and mutant alleles of *SMO4* and *MTR4*, which participate in 5.8S rRNA maturation (Lange et al., 2011; Micol-Ponce et al., 2020) and *RRP7* and *PARALLEL1* (*PARL1*; also named *NUCLEOLIN1*), which act in 18S rRNA maturation (Pontvianne et al., 2007; Micol-Ponce et al., 2018).

All of the double mutants showed synergistic phenotypes (Figure 5). Leaves of *rps24b-*2 *smo4-2* plants exhibited increased serration and took on a twisted appearance into a spiral (Figure 5F). *rps24b-*2 *mtr4-2* plants exhibited delayed growth, lanceolate leaves and strong drops in rosette size and plant height (Figure 5G). *rps24b-2 parl1-2* plants strongly accumulated anthocyanins, exhibiting dark, serrated, pointed leaves and small, compact rosettes (Figure 5H). The *rps24b-2 mtr4-2* and *rps24b-2 parl1-2* double mutants completed their life cycles but had poor fertility (Figure 5I, J, M-P). We sowed 108 F_2_ seeds from an *rps24b-2 × rrp7-1* cross and isolated two double mutants, which exhibited very strong mutant phenotypes (Supplemental Figure 7). Only one plant completed its life cycle and produced 20 F_3_ seeds, only one of which germinated, but the seedling died a few days later, suggesting early postembryonic lethality of the *rps24b-2 rrp7-1* genotype.

**Figure 5.**
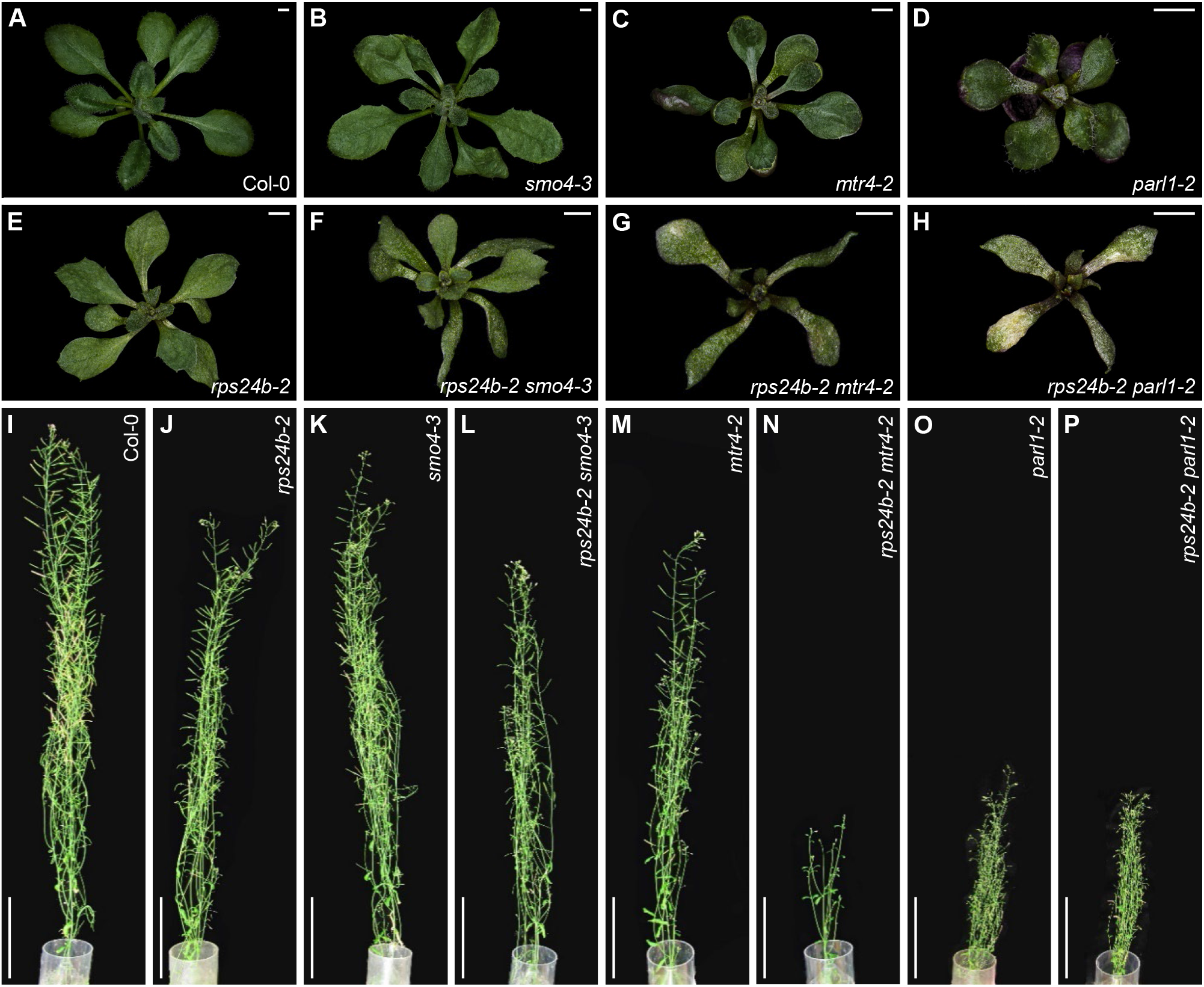
Genetic interactions between *rps24b-2* and mutant alleles of genes encoding RBFs. Rosettes of Col-0 (A), *smo4-3* (B), *mtr4-2* (C), *parl1-2* (D), *rps24b-2* (E), *rps24b-2 smo4-3* (F), *rps24b-2 mtr4-2* (G) and *rps24b-2 parl1-2* plants (H). From left to right, adult plants of Col-0 (I), *rps24b-2* (J), *smo4-3* (K), *rps24b-2 smo4-3* (L), *mtr4-2* (M), *rps24b-2 mtr4-2* (N), *parl1-2* (O) and *rps24b-2 parl1-2* (P). Photographs were taken 21 das (A-H) or 57 das (I-P). Scale bars, 2mm (A-H), or 5 cm (I-P).

We analyzed 45S pre-rRNA maturation in the viable double mutants, finding similar levels of P-A_3_ or P′-A_3_/18S-A_3_/18S-A_2_ pre-rRNAs in *rps24b-2 smo4-3*, *rps24b-2 mtr4-2* and *rps24b-2 parl1-2* compared to the single mutants using the S2 and S7 probes (Figure 6A and B). The *smo4-3* and *mtr4-2* single mutants accumulate 7S and 5.8S+10 pre-rRNAs, respectively (Lange et al., 2011; Micol-Ponce et al., 2020). Unexpectedly, using the S9 probe, we detected higher levels of 7S pre-rRNA in *rps24b-*2 *smo4-3* and of 5.8S+10 pre-rRNA in *rps24b-2 mtr4-2* compared to the *smo4-3* and *mtr4-2* single mutants, even though *rps24b-2* did not accumulate either of these pre-rRNA species (Figure 6C and D).

**Figure 6.**
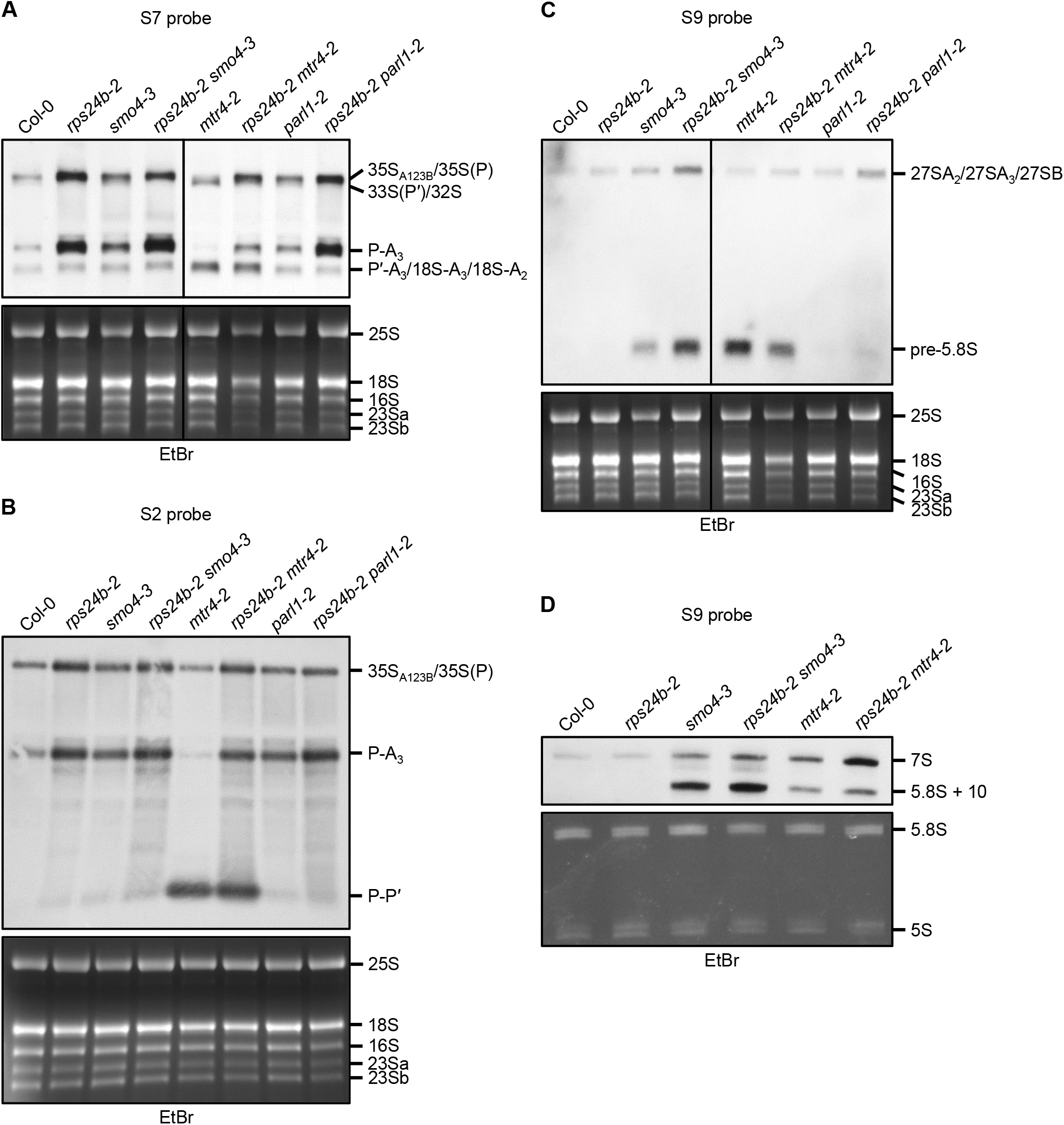
rRNA maturation in double mutants between *rps24b-2* and alleles of genes encoding RBFs. (A-D) Visualization of the processing of the 5.8S, 18S and 25S rRNA precursors using gel blot analysis. Total RNA from seedlings collected 15 das was separated on (A-C) formaldehyde-agarose or (D) polyacrylamide-urea gels, transferred to a nylon membrane and hybridized with the S7 (A), S2 (B), or S9 probe (C and D). EtBr, ethidium bromide-stained gels, visualized before blotting, serving as loading controls. Similar results were obtained in at least two independent experiments.

To assess the subcellular localization of 5.8S pre-rRNA in the *rps24b-*2 *smo4-3* and *rps24b-*2 *mtr4-2* double mutants, we performed an RNA-FISH assay. We used the 23-nt long S9 probe, which hybridizes to the ITS2 region adjacent to the 5.8S rRNA coding sequence, and allows detection of all 5.8S pre-rRNA, but not mature 5.8S rRNA (Figure 4A). In agreement with the results of RNA gel blot analysis (Figure 6C and D), we observed very low fluorescence in the nucleoli of wild-type Col-0 and L*er*, higher fluorescence in the *api6*, *rps24a-1*, *rps24b-2*, *smo4-3* and *mtr4-2* single mutants, and much higher fluorescence in the *rps24b-*2 *smo4-3* and *rps24b-*2 *mtr4-2* double mutants (Supplemental Figure 8). The enhanced fluorescence that we observed in the nucleoli of *rps24* mutants compared to Col-0 and L*er* is likely due to the accumulation of the 35S_A123B_/35S(P) species that we detected in RNA gel blots using the S7 and S9 probes (Figure 4B-D).

In the RNA-FISH assays, we also observed enlarged nucleoli in *rps24b-2* and (especially) *smo4-3 rps24b-2* and *mtr4-2 rps24b-2* double mutant plants compared to Col-0 (Supplemental Figure 8). Perturbations in the size and organization of the nucleolus are hallmarks of nucleolar stress, which occurs when ribosome biogenesis is defective, as previously observed in the nucleoli of *smo4* plants (Micol-Ponce et al., 2020). Therefore, we measured the areas of the nucleolus and nucleoplasm using the fluorescence of the S9 probe and of nuclei stained with DAPI. Both the nucleus and nucleolus of *api6* and *rps24b-2* (but not *rps24a-1*) were larger than those of the respective wild types (Supplemental Figure 8A2 and A3). Moreover, the nucleus and nucleolus were greatly enlarged in both the *smo4-3 rps24b-2* and *mtr4-2 rps24b-2* double mutants (Supplemental Figure 8D-F, J-L, P-R, V-A3).

To determine whether the organization of the nucleolus was perturbed in the *rps24* mutants, we used an antibody against the nucleolar marker fibrillarin to examine the roots of Col-0, *rps24a-1* and *rps24b-2* seedlings. We did not observe differences in nucleolar organization between the wild type and the mutants (Supplemental Figure 9), suggesting that the defective 45S pre-rRNA processing in the *rps24* mutants causes the enlargement but not disorganization of the nucleolus.

Taken together, these results suggest that RPS24B participates (directly or indirectly) in the maturation of 5.8S rRNA. Since the *rps24a-1* and *rps24b-2* single mutants do not accumulate any 5.8S pre-rRNA, RPS24B might play a lesser role in 5.8S maturation compared to SMO4 and MTR4; therefore, the absence of RPS24B is only perceived in the absence of SMO4 and MTR4. Another interpretation is that RPS24 participates in the repression of 45S rDNA transcription. Under this second scenario, the accumulation of 5.8S pre-rRNAs would be more evident in the *rps24b* background than in the *smo4-3* and *mtr4-2* single mutant backgrounds due to the presence of more 5.8S pre-rRNAs to process.

### 45S pre-rRNAs accumulate in the *rps24* mutants

Kinetic analysis of Arabidopsis 45S pre-rRNA processing has revealed three rate-limiting steps: the processing of 35S, 32S, and P-A_3_ pre-rRNAs; 32S and P-A_3_ pre-rRNAs are the first two precursors in the two alternative pathways, the 5′-ETS first pathway and the ITS1-first pathway, respectively (Supplemental Figure 1). The level of 35S pre-rRNA is the major indicator of the transcription rate of 45S rDNA (Shanmugam et al., 2021). Our results show that a reduction in RPS24 activity lower the rate of 45S pre-rRNA processing, as shown by the accumulation of pre-rRNAs produced by two of the rate-limiting steps, 35S_A123B_/35S(P) and P-A_3_. Our results also suggest that 45S rDNA transcription is upregulated in the *rps24* mutants.

Therefore, we performed RT-PCR to investigate the presence of the four different 45S rDNA variants (*VAR*s), which differ in their 3′-ETS (Figure 8A), using the p3+p4 primer pair (Figure 7A and B; Supplemental Table 2). We did not detect any differences in the intensities of the PCR bands using genomic DNA as a template (Figure 7C). However, the intensities of the PCR bands corresponding to *VAR4*, *VAR2* and *VAR3* were higher in *rps24a-1*, and those corresponding to all four *VARs* were higher in *rps24b-2*, compared to the wild type (Figure 7D). Since changes in the relative abundance of 45S rRNA variants are usually interpreted as changes in 45S rDNA transcription (Kojima et al., 2007; Pontvianne et al., 2010; Durut et al., 2014), these results suggest that RPS24B represses 45S rDNA expression.

**Figure 7.**
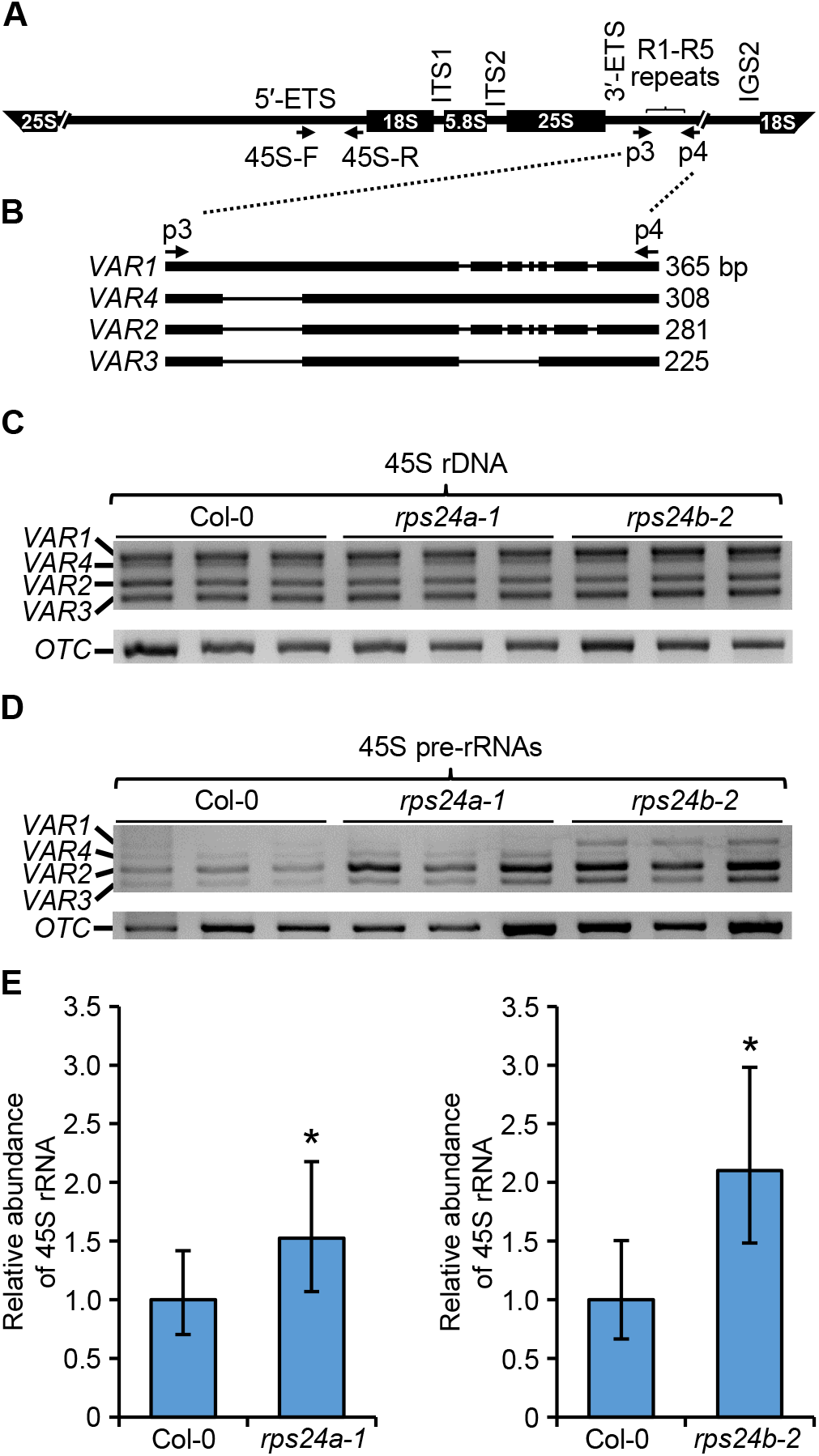
45S rDNA expression in the *rps24a-1* and *rps24b-2* mutants. (A and B) Schematic representation of 45S pre-rRNA (A) and its 3′-ETS polymorphic region (B), modified from Micol-Ponce *et al*. (2018). (C and D) PCR analysis of the relative abundance of 45S rDNA variants using genomic DNA (C) and cDNA templates (D) from three biological replicates obtained from Col-0, *rps24a-1* and *rps24b-2* plants. The analysis was performed twice with similar results. (E) RT-qPCR analysis of 45S rRNA abundance. Eight biological replicates were performed per genotype (Col-0, *rps24a-1*, and *rps24b-2*), with three technical replicates in each experiment, resulting in 24 amplifications per genotype. The analyses were carried out using ExpressionSuite Software. Asterisks indicate a significant difference with Col-0 in a Mann-Whitney *U* test (n = 8) (\**P* < 0.05).

To confirm these results, we carried out RT-qPCR analysis using the 45S pre-rRNA-45S-F and 45S pre-rRNA-45S-R primers, which hybridize to the 5′-ETS (Supplemental Table 2; Zhu et al., 2016). We detected higher levels of 45S pre-rRNA in the *rps24* mutants than in wild-type plants (Figure 7E).

### RPS24B-GFP co-precipitates with nucleolar factors involved in ribosome biogenesis and pre-mRNA splicing

To identify nucleolar interactors of RPS24B, we carried out co-immunoprecipitation (Co-IP) assays with an anti-GFP antibody in Col-0 plants harboring the *35S_pro_:RPS24B:GFP* transgene. The raw list of 2,635 proteins that co-purified with RPS24B-GFP was filtered by considering only those classified as high-confidence proteins (q-value <0.01) with at least two peptide spectra matches (PSMs) in each of the three biological replicates (Supplemental Dataset 2). Several additional filters were used, which discarded proteins that co-immunoprecipitated with all four other proteins examined (none related to ribosome biogenesis), as we considered these to be spurious interactions. We analyzed the predicted suborganellar localizations of the interactors using MuLOCDeep (Jiang et al., 2021), discarding those that were not predicted to be nucleolar proteins, which reduced the list of putative interactors to 25 proteins (Supplemental Dataset 2). We refined the annotations of these proteins using the TAIR or Aramemnon database and those identified in two independent nucleolar proteomic analyses (Palm et al., 2016; Montacié et al., 2017). In addition, we added the pathways that were enriched in genes that were co-expressed with the genes encoding the 25 putative interactors of RPS24B obtained from the ATTEDII database (Obayashi et al., 2022).

We found ten proteins encoded by genes whose co-expression was enriched in ribosome components, which included six RPs, six whose co-expression was enriched in genes involved in ribosome biogenesis, three in spliceosome components, and two in RNA transport, but that were components of the spliceosome: PRE-MRNA PROCESSING FACTOR 8 (PRP8), the main component of the spliceosome, and one of its major interactors during pre-mRNA splicing, the putative RNA helicase BAD RESPONSE TO REFRIGERATION2 (BRR2; Nguyen et al., 2013).

Among the putative interactors identified by co-immunoprecipitation that were encoded by genes that are co-expressed with genes involved in ribosome biogenesis, we found AT3G05060 (*NOP58-2*) and AT5G27120 (*NOP58-1*); these genes encode the co-orthologs of *NOP58* in yeast and humans, which are putative components of the C/D small nucleolar ribonucleoprotein (snoRNP) complex. AT1G31970 and AT1G77030, encoding DEA(D/H)-box RNA helicases, were two other co-expressed genes that are putative orthologs of yeast *DEAD-box protein 3* (*DBP3*) and *DBP10*, respectively, both of which are involved in 60S subunit biogenesis. DBP3 functions in the endonucleolytic cleavage of site A_3_ in ITS1, upstream of 5.8S rRNA (Weaver et al., 1997), and DBP10 functions in the cleavage of sites C_1_ and C_2_ in the ITS2 (Burger et al., 2000). AT1G31970 encodes STRESS RESPONSE SUPPRESSOR 1 (STRS1), a nucleolar- and chromocenter-localized protein that undergo stress-mediated relocalization and is involved in epigenetic gene silencing at heterochromatic loci; its mutants show reduced DNA methylation at these loci (Khan et al., 2014). AT1G77030 encodes the putative DEAD-box ATP-dependent RNA HELICASE 29 (RH29), which has not been studied in Arabidopsis. AT3G56510, AT5G04600 and AT5G57120 have not been studied either. AT3G56510 is annotated at the Aramemnon database (Schwacke et al., 2003) as the putative ortholog of yeast Eighteen S rRNA Factor 2 (Esf2), a nucleolar component of the small subunit processome (SSU) involved in early steps of 18S rRNA maturation. Yeast Esf2 associates with 5′-ETS and is required for SSU assembly and compaction. The *esf2* mutants show defective cleavage of sites A0 (within the 5′-ETS) through A2 (within the ITS1), therefore exhibiting reduced 18S rRNA production (Hoang et al., 2005). AT5G04600 encodes the putative ortholog of human NUCLEOLAR PROTEIN INTERACTING WITH THE FHA DOMAIN OF MKI67 (NIFK), an RBF involved in the endonucleolytic ITS2 cleavage that is required for 28S and 5.8S rRNA maturation (Pan et al., 2015). Finally, AT5G57120 encodes the putative ortholog of yeast serine rich protein 40 (SRP40) and human nucleolar and coiled-body phosphoprotein 1 (NOLC1), also named 140 kDa nucleolar phosphoprotein (Nopp140), which might function as chaperones of snoRNP complexes (Yang et al., 2000).

## DISCUSSION

### The Arabidopsis *RPS24A* and *RPS24B* paralogs exhibit combined haploinsufficiency

The ribosome has traditionally been considered to be a highly conserved housekeeping machinery, as it is present in all cells of all organisms. In prokaryotic and eukaryotic ribosomes, the number of RPs is almost identical, i.e., 55 and 79, respectively. However, proteomic analyses of animals, plants and fungi have revealed the existence of different ribosomes with heterogeneous compositions and stoichiometry, suggesting that specialized ribosomes might translate different sets of mRNAs. Such heterogeneity in composition results from the existence of different paralogous genes produced by gene duplications. These genes, with similar architectures and variable numbers among species, encode quasi-identical RPs [reviewed in Martinez-Seidel et al. (2020; Petibon et al. (2021)].

Plants have undergone more frequent gene and genomic duplications during evolution compared to metazoans, leading to families of RPs with two to seven paralogs in Arabidopsis (Carroll et al., 2008). In Arabidopsis, the RPs of the cytoplasmic ribosome appear to be encoded by 234 functional genes, whereas the ribosomes of humans and yeast include proteins encoded by 85 and 137 genes, respectively (Barakat et al., 2001; Chang et al., 2005; Carroll et al., 2008). In the quasi-identical RP pairs studied to date, one of the two paralogous genes is sometimes expressed at a higher level and contributes more to ribosome structure than the other gene. In other cases, their levels of expression and contributions to the ribosome are equivalent [reviewed in Petibon et al. (2021)]. Loss of function of either gene of a pair produces a similar mutant phenotype, indicating that both paralogs are required for normal development (Savada and Bonham-Smith, 2014; Salih et al., 2020; Xiong et al., 2021).

Haploinsufficiency describes the requirement for more than one wild-type allele for normal development or viability for single-copy genes (Meinke, 2013). Combined haploinsufficiency describes a situation in which two paralogous genes behave as a single haploinsufficient unit with four alleles (Hawley and Gilliland, 2006). *RPS24A* and *RPS24B* provide such an example of combined haploinsufficiency, since three wild-type doses of *RPS24* are required for proper plant development and two for viability. It is worth noting that the single-copy human *RPS24* gene is a haploinsufficient locus. Haploinsufficiency is observed in several human genetic diseases, such as Diamond-Blackfan anemia (DBA), a ribosomopathy (Gazda et al., 2006) associated with the loss-of-function of *RPS24*, as well as other genes encoding RPs (Tyagi et al., 2020). Ribosomopathies are diseases caused by mutations in genes encoding components of the ribosome, with tissue-specific effects and an increased risk of cancer [reviewed in Kang et al. (2021)].

*RPS24A* and *RPS24B* have similar expression patterns and contributions to the composition of Arabidopsis ribosomes, pointing to their functional equivalence and partial redundancy. However, the mRNA levels of *RPS24A* and *RPS24B* are lower than those of other genes encoding RPs (Savada and Bonham-Smith, 2014). Here we demonstrated that these genes do not show dosage compensation; i.e., *RPS24A* is not more expressed in the absence of *RPS24B*. A proper stoichiometry of ribosome structural components is crucial for ribosome function (Warner, 1999; Slavov et al., 2015; Ni and Buszczak, 2023), and our results suggest that the amount of RPS24A and RPS24B produced in Arabidopsis cells is limiting for ribosome production. Indeed, reduction in the wild-type doses of either protein in *rps24a* and *rps24b* single mutants and diheterozygous plants (Figures 1 and 2) decreases the number of productive ribosomes, resulting in an inadequate rate of protein synthesis. This phenomenon of non-allelic non-complementation is also observed in mutants affected in other Arabidopsis genes encoding RPs, including *RPS6*, *RPL5*, *RPL23* and *RPL36* (Degenhardt and Bonham-Smith, 2008; Fujikura et al., 2009; Creff et al., 2010; Casanova-Sáez et al., 2014).

### RPS24A and RPS24B function as RBFs during 18S rRNA maturation, like their yeast and human orthologs

Gene duplications have allowed some RPs to acquire additional functions that might or might not be related to translation. In Arabidopsis, besides their functions as structural proteins of the ribosome, only RPS2 was shown to function as an RBF during the maturation of 25S and 18S rRNA, and RPS6 in the transcriptional regulation of 45S rDNA (Kim et al., 2014; Hang et al., 2021). PROTEIN ARGININE METHYLTRANSFERASE 3 (PRMT3) and RPS2B (one of the six paralogs encoded in the Arabidopsis genome) facilitate the assembly and disassembly of the SSU processome and repress nucleolar stress (Hang et al., 2021). RPS6, encoded by two functionally equivalent paralogs, interacts with HISTONE DEACETYLASE 2 (HDA2) in the nucleus and nucleolus and binds to the promoter of 45S rDNA, controlling its transcription (Kim et al., 2014). Furthermore, RPL10 is related to responses to ultraviolet light stress and viral infection, and RPL24 is involved in microRNA biogenesis (Carvalho et al., 2008; Ferreyra et al., 2010a; Ferreyra et al., 2010b; Li et al., 2017).

Our results show that both Arabidopsis RPS24A and RPS24B function as RBFs in the early steps of 45S pre-rRNA processing, specifically at the cleavage of the P′ site of the 5′-ETS. In the *rps24b-2*, *api6* and *rps24a-1* mutants, we detected the accumulation of 35S_A123B_/35S(P) pre-rRNA (the first intermediate in 45S pre-rRNA processing) and a reduction in the levels of 33S(P′)/32S species (the second intermediate in this processing; Supplemental Figure 1). We also detected defects in the processing of 18S pre-rRNA: the *rps24b-2*, *api6* and *rps24a-1* mutants accumulated P-A_3_ pre-rRNA and showed reduced levels of P′-A_3_/18S-A_3_/18S-A_2_ pre-rRNAs (Figure 4). These results suggest that the loss-of-function of *RPS24A* and *RPS24B* globally reduces the processing of 5′-ETS, as occurs in response to the loss-of-function of yeast and human *RPS24* (Ferreira-Cerca et al., 2005; Choesmel et al., 2008; Robledo et al., 2008). Moreover, the *rps24b* mutant showed an imbalance in the 25S/18S ratio, suggesting that alterations in the processing of pre-rRNAs of the ITS1-first pathway lead to a decrease in 18S rRNA levels.

### *RPS24B* genetically interacts with *NOP53* (*SMO4*) and *MTR4*

We obtained double mutant combinations of *rps24b-2* with mutant alleles of genes involved in different steps of 45S pre-rRNA processing. Two such double mutants exhibited synergistic phenotypes: *rps24b-2 rrp7-1* was embryonic lethal, and *rps24b-2 mtr4-2* was dwarf, with poor fertility. The phenotype of *rps24b-2 smo4-3* was also synergistic, as this mutant was also nearly fully sterile (Figure 5).

We previously demonstrated that RRP7 is involved in 18S rRNA maturation and acts as a repressor of 45S rDNA transcription. The *rrp7* mutant has a reduced 18S/25S rRNA ratio and accumulated P-A_3_ pre-rRNA (Micol-Ponce et al., 2018), similar to the *rps24* mutant. Moreover, we reasoned that RRP7 might function as a repressor of 45S rDNA, as *rrp7* showed heterochronic expression of *VAR1* and increased *VAR3* expression (Micol-Ponce et al., 2018). Here, we determined that *rps24a-1* and *rps24b-2* contained increased levels of 45S pre-rRNA (Figure 7), which might explain the synergistic phenotype of the *rps24b-2 rrp7-1* double mutant. However, due to the lethality of *rps24b-2 rrp7-1*, we were unable to examine the effects of the combination of both mutations on the maturation of 18S rRNA and/or the transcription of 45S rDNA.

NOP53 (SMO4 in Arabidopsis) and MTR4 are RBFs that function in the maturation of 5.8S rRNA (Lange et al., 2011; Micol-Ponce et al., 2020). MTR4 is a cofactor of the exosome, which also functions in the degradation of P-P′, the by-product of 5′-ETS (Lange et al., 2011). Loss-of-function of *SMO4* or *MTR4* leads to the accumulation of 7S or 5.8S + 70-nt pre-rRNAs, respectively (which are precursors of 5.8S rRNA that are processed in the nucleolus), but not 6S rRNA (the final precursor of 5.8S rRNA maturation that is processed after its export to the cytoplasm; Lange et al., 2011; Micol-Ponce et al., 2020). In the current study, we did not detect the accumulation of P-P′ or any 5.8S pre-rRNA in the *rps24a* or *rps24b* mutants, suggesting that RPS24A and RPS24B do not participate in these steps of 45S pre-rRNA processing. However, we unexpectedly detected the over-accumulation of specific 5.8S pre-rRNAs in *smo4-3* and *mtr4-2* and in the double mutants *rps24b-2 smo4-3* and *rps24b-2 mtr4-2* (Figure 6). Such over-accumulation may explain their synergistic phenotypic defects. Our RNA-FISH analysis with the S9 probe confirmed these results, revealing strong fluorescent signals in the nucleoli of the double mutants (Supplemental Figure 8).

One possible explanation of the above-mentioned results is that RPS24B (and possibly RPS24A) functions in 5.8S rRNA maturation, albeit to a lesser extent than SMO4 and MTR4. An alternative hypothesis is that the over-accumulation of 7S and 5.8S + 70-nt pre-rRNAs, which we detected in *rps24b-2 mtr4-2* and *rps24b-2 smo4-*3 plants, respectively, is due to an increase in rDNA transcription caused by the *rps24b-2* mutation. If more pre-rRNAs are available for processing, the steps involving MTR4 and SMO4 are likely to be more strongly affected in the double mutants. We consider this second hypothesis to be more plausible, as it also might explain the synergistic phenotype of the *rps24b-2 rrp7-1* and *rps24b-2 parl1-2* double mutants.

### The predominant nucleolar localization of RPS24B and its putative interactors also support its role as an RBF

Our results using the RPS24B-GFP fusion protein indicate that RPS24B is primarily located in the nucleolus, with some presence in the cytoplasm; these observations are consistent with previous predictions and proteomic analyses. This distribution reflects the dual role of RPS24B as both a structural protein of the ribosome and an RBF involved in early steps of ribosome biogenesis and the transcriptional repression of 45S rDNA.

Using different filters that included sub-organellar localization, we identified 25 nucleolar proteins that co-immunoprecipitated with RPS24B-GFP, which we considered to be its most likely interactors. Among these were RBFs involved in the synthesis of the 60S and 40S subunits, including STRS1 (AT1G31970), AT1G77030 and AT3G56510, which are putative orthologs of yeast DBP3, DBP10 and ESF2, respectively, putative ortholog of yeast, as well as NIFK. We speculate that STRS1 might also be involved in repressing 45S rDNA, which would be difficult in the *rps24* background if RPS24A, RPS24B, or both proteins facilitate access to the epigenetic silencing machinery at the promoter of 45S rDNA. In fact, STRS1 colocalized with HDA2 in the nucleolus (Khan et al., 2014). Like RPS6, another RP that plays extra-ribosomal roles in the maturation of 18S rRNA and the transcriptional regulation of 45S rDNA (Kim et al., 2014), RPS24A and RPS24B might also act in this capacity.

We also identified AT5G57120 as an RPS24B-GFP interactor. AT5G57120 is the putative ortholog of human NOLC1, which colocalizes with RNA Pol I in the nucleolus at 47S rDNA and might activate its transcription (Chen et al., 1999). The interaction between the protein encoded by AT5G57120 and RPS24B might inhibit access of the-encoded protein to the 45S rDNA promoter. RPS24B, and probably RPS24A, might also act as chaperones to facilitate the incorporation of RBFs into their corresponding pre-rRNA substrates, as observed for other RPs. Although we did not find SMO4 or MTR4 among the RPS24B-GFP interactors, we cannot exclude the possibility that RPS24B and RPS24A function in the maturation of 5.8S rRNA. However, we consider this possibility unlikely, as previously argued. Interestingly, we also identified two well-known splicing factors among the proteins that co-immunoprecipitated with RPS24B-GFP: PRP8 and the RNA helicase BRR2. PRP8 was previously identified in a U3 snoRNA ribonucleoprotein complex isolated from cauliflower (*Brassica oleracea*) inflorescences (Samaha et al., 2010) and in a proteomic analysis of the Arabidopsis nucleolus (Palm et al., 2016). We cannot exclude the possibility that PRP8 and BRR2 function in ribosome biogenesis, as the expression of *RPS24B-GFP* was unable to rescue the mutant phenotype of *rps24b-2* plants, suggesting that many RNA helicases act in more than one pathway. Indeed, yeast Prp43 and its human ortholog DEAD-box helicase 5 (DDX5) are multifunctional RNA helicases that participate in ribosome biogenesis and pre-mRNA splicing, among other RNA metabolic steps [reviewed in Bohnsack et al. (2022)]. However, our assay may have generated false positives, since RPS24B-GFP was unable to rescue the mutant phenotype of *rps24b-2* plants.

## MATERIALS AND METHODS

### Plant material and growth conditions

The wild-type Arabidopsis (*Arabidopsis thaliana*) (L) Heynh. Columbia-0 (Col-0) and Landsberg *erecta* (L*er*) accessions were provided by the Nottingham Arabidopsis Stock Centre (NASC) and propagated in our laboratory. The *api6* mutant in the L*er* background was isolated in the laboratory of J.L. Micol after ethyl methanesulfonate (EMS) mutagenesis (Berná et al., 1999). Seeds of *rps24b-2* (SALK_000766; Wang et al., 2018), *rps24a-1* (SALK_126799), *smo4-3* (SALK_071764; Micol-Ponce et al., 2018), *mtr4-2* (SAIL_50_C11; Lange et al., 2011), *rrp7-1* (SAIL_628_F08; Micol-Ponce et al., 2018) and *parl1-2* (SALK_002764; Petricka and Nelson, 2007) were also provided by NASC. Seed sterilization, sowing, plant culture and crosses were performed as previously described (Berná et al., 1999; Ponce et al., 1999).

### Positional cloning of the *api6* mutation and genotyping of single and double mutants

Genomic DNA extraction and mapping of the *api6* mutation were performed as previously described (Ponce et al., 1999; Ponce et al., 2006). The primers used for fine-mapping are listed in Supplemental Table 1. The *api6* point mutation was identified by Sanger sequencing using the primers listed in Supplemental Table 2. The presence of T-DNA insertions in *RPS24A*, *RPS24B*, *SMO4*, *RRP7, MTR4* and *PARL1* was verified by PCR using the primers shown in Supplemental Table 2.

Most Sanger sequencing reactions and electrophoreses were carried out in our laboratory with ABI PRISM BigDye Terminator Cycle Sequencing kits and an ABI PRISM 3130xl Genetic Analyzer (Applied Biosystems). Some Sanger sequencing reactions were carried out by Stab Vida (Portugal).

### Construction of transgenic lines

The constructs used in this study were generated by Gateway cloning as described in Sánchez-García et al. (2015) using the pGEM-T Easy221 entry vector and the pMDC83 destination vector (Curtis and Grossniklaus, 2003). To generate the *35S_pro_:RPS24B:GFP* transgene, the full-length coding region (without the stop codon) of *RPS24B* was amplified by PCR using genomic DNA from Col-0 as a template and primers that included the *att*B1 and *att*B2 sequences (Supplemental Table 2). The integrity of the constructs was verified by Sanger sequencing. Chemically competent *Escherichia coli* DH5α cells were transformed with the Gateway cloning products by the heat shock method, and *Agrobacterium tumefaciens* C58C1 cells were transformed by electroporation with the individual destination vectors carrying each insert of interest.

### Plant morphological and histochemical analyses

The rosettes of plants were photographed under a Nikon SMZ1500 stereomicroscope equipped with a Nikon DS-Ri2 digital camera. For large rosettes, several photographs from the same plant were assembled with the Photomerge tool of Adobe Photoshop CS3 software (Adobe). The backgrounds of the photographs were homogenized using Adobe Photoshop CS3 software without modifying the rosettes.

Fluorescence and confocal laser-scanning microscopy images were taken under a Leica STELLARIS microscope and processed using Leica Application Suite X (LAS X) software. For nuclear staining, the roots of seedlings collected 5 days after stratification (das) were immersed in 0.5 µg/mL 4′,6-diamidino-2-phenylindole (DAPI) for 5 min and washed two times with water. DAPI, GFP, tetramethylrhodamine-5-isothiocyanate (TRITC) and cyanine 3 (Cy3) were excited at 405, 488, 503 and 506 nm, respectively, and their emissions collected at 425/727, 515/730, 530/732 and 511/726 nm, respectively.

### RT-PCR and RNA gel-blot analysis

Genomic DNA was extracted using a DNeasy Plant mini kit (Qiagen) from three biological replicates per genotype (leaves from a pool of three seedlings collected 15 das from three different Petri plates). Total RNA was extracted using TRIzol Reagent (Invitrogen) for RT-PCR and semiquantitative RT-PCR and an RNeasy Plant Mini Kit (Qiagen) for RT-qPCR RNA gel blots. Three biological replicates per genotype were used for RT-PCR and semiquantitative RT-PCR, and eight were used for RT-qPCR. Each biological replicate included the aerial tissues from a pool of three seedlings collected 15 das from three different Petri plates. In all RT-PCR assays, RNA was treated twice with Turbo DNase (Invitrogen), and reverse transcription was carried out using random hexamer primers and Maxima Reverse Transcriptase (Invitrogen). The *ORNITHINE CARBAMOYLTRANSFERASE* (*OTC*) and *ACTIN2* (*ACT2*) housekeeping genes were used as internal controls: *OTC* for RT-PCR and semiquantitative RT-PCR, and *ACT2* for RT-qPCR assays. Primers for all RT-PCR assays and those used as controls for genomic DNA amplification in the analysis of 45S rDNA variant expression are listed in Supplemental Table 2.

For RNA gel-blot analysis, four digoxigenin (DIG) labeled probes were used, whose sequences were obtained from Lange et al. (2011); the S6, S7 and S9 probes were oligonucleotides labeled at both the 3′ and 5′ ends, synthetized by Sigma-Aldrich. The S2 probe was synthetized by PCR using genomic DNA as a template, DIG-11-dUTP (Roche) and the S2-Fw and S2-Rv primers (Supplemental Table 2). Two µg of total RNA per sample were loaded onto 1.2% (w/v) agarose and 2.12% (v/v) formaldehyde or 6% (w/v) polyacrylamide gels. Electrophoresis, hybridization and detection were performed as described in Micol-Ponce et al. (2018; Micol-Ponce et al. (2020).

### RNA-FISH and immunolocalization

RNA fluorescence *in situ* hybridization (RNA-FISH) was performed as described by Parry et al. (2006) and modified in Micol-Ponce et al. (2020) using the S9 probe labeled with Cy3 dye (Eurofins Genomics) at a concentration of 0.5 µg/mL. Leaves were mounted onto slides with Vectashield antifade mounting medium (Vector Laboratories) containing 0.01 µg/mL DAPI, which was used as a nuclear marker.

Fibrillarin immunolocalization was performed following Pasternak et al. (2015), as modified by Micol-Ponce et al. (2020). In brief, roots of seedlings were collected at 5 das and fixed for 40 min at 37°C in a solution containing 2% (w/v) paraformaldehyde in 1× microtubule-stabilizing buffer (50 mM PIPES, 5 mM MgSO_4_, and 5 mM EGTA, pH 6.9) and 0.1% (v/v) Triton X-100. An anti-fibrillarin [38F3] (Abcam) primary antibody was used at a 1:250 dilution, and a TRITC-conjugated anti-mouse IgG (Sigma Aldrich) secondary antibody was used at a 1:500 dilution. Nuclei were stained for 10 min with 0.2 µg/mL DAPI and washed for 5 min before mounting the samples on slides with water. Nuclei and nucleoli areas were measured as described in Micol-Ponce et al. (2020) using the NIS Elements AR3.1 (Nikon) image-analysis package.

### Co-immunoprecipitation assay

For co-immunoprecipitation assays, three biological replicates were used, each consisting in 1 g of rosettes of *35S_pro_:RPS24B:GFP* transgenic plants in the Col-0 background collected 15 das. The tissue was manually ground and proteins were extracted as described in Navarro-Quiles et al. (2022). Co-immunoprecipitation was carried out using a µMACS GFP Isolation kit (Milteny Biotec). The effectiveness of the immunoprecipitation of RPS24B-GFP was checked by immunoblotting using anti-GFP-HRP antibody (Milteny Biotec).

The co-immunoprecipitated proteins were identified at the Centro Nacional de Biotecnología (CNB) Proteomics facility (Madrid, Spain) by liquid chromatography followed by tandem mass spectrometry (LC-MS/MS) using an Orbitrap Exploris 240 mass spectrometer. Tandem mass peptide spectra were searched against Arabidopsis protein sequences in the Araport11 database using the MASCOT search engine (Matrix Science). Peptide sequences identified with a false discovery rate (FDR) < 1% were considered to be valid. To increase the confidence of the results, only the proteins identified with at least two peptides in each of the three biological replicates were considered for analysis.

### Accession numbers

Sequence data from this article can be found at The Arabidopsis Information Resource (https://www.arabidopsis.org/) under the following accession numbers: *RPS24A* (AT3G04920), *RPS24B* (AT5G28060), *RRP7* (AT5G38720), *SMO4* (AT2G40430), *MTR4* (AT1G59760), and *PARL1* (AT1G48920). Although AT3G04920 and AT5G28060, which encode RPS24A and RPS24B, have been renamed as ES24Z and ES24Y, respectively, according to a new nomenclature for RPs (Scarpin et al., 2022), we used their old names in this work, since the *rps24b-2* allele of *RPS24* were previously studied (Horiguchi et al., 2011).

## ACKNOWLEDGEMENTS

The authors thank F. Lozano, J. Castelló, D. Navarro and M. Gomariz for their excellent technical assistance, and J.L. Micol for useful discussions, comments on the manuscript, and providing the *api6* mutant, as well as for the use of his facilities.

## AUTHOR CONTRIBUTIONS

M.R.P. obtained funding and conceived, designed and supervised research; all authors performed research and wrote the paper.

## SUPPLEMENTAL FIGURE LEGENDS

**Supplemental Figure 1.** Overview of 45S pre-rRNA processing in Arabidopsis. Colored boxes represent the sequences of the three mature rRNAs transcribed from the 45S rDNA genes. Vertical arrows indicate endonucleolytic cleavages; letters indicate the cleavage site in the corresponding pre-rRNA. Only the relevant factors and the human (H) and yeast (Y) 18S pre-rRNAs relevant to this study are represented. Based on information from Sáez-Vasquez and Delseny (2019).

**Supplemental Figure 2.** Sequence conservation between the RPS24A and RPS24B paralogs. Amino acid sequence alignment of Arabidopsis RPS24B and RPS24A. The 14 amino acids that are absent from the RPS24B variant produced by the expression of the *api6* allele are highlighted in red letters. Identical and similar residues are shaded in black and gray, respectively. Asterisks and dots in the consensus line indicate identical and conserved residues, respectively. Numbers indicate residue positions. The alignment was obtained using ClustalW2 and shaded with Boxshade 3.21 (http://www.ch.embnet.org/software/BOX_form.html). Nuclear and nucleolar localization sequences predicted with LOCALIZER (https://localizer.csiro.au/) and NoD software (http://www.compbio.dundee.ac.uk/www-nod/index.jsp) are underlined in black and green, respectively.

**Supplemental Figure 3.** R*P*S24B expression analysis. (A) Schematic representation of *RPS24B* and its mutant alleles used in this study. Gene structure is represented as described in the legend of Figure 1. Black arrows represent the primers used for semiquantitative RT-PCR amplifications in (B). (B) Semiquantitative RT-PCR analysis of *RPS24B* in the *rps24a* and *rps24b* mutants used in this study. The bands for each PCR were visualized after 25, 30 and 35 cycles of amplification. Total RNA was extracted from seedlings collected at 15 das. Transcripts from the *OTC* gene were used as an internal control. The primer sequences used are shown in Supplemental Table 2.

**Supplemental Figure 4.** Predicted localization of RPS24A. (A-B) Predicted subcellular (A) and sub-organellar (B) localization of RPS24A using MULocDeep software (https://mu-loc.org/).

**Supplemental Figure 5.** Predicted localization of RPS24B. (A-B) Predicted subcellular (A) and sub-organellar (B) localization of RPS24B protein using MULocDeep software (https://mu-loc.org/).

**Supplemental Figure 6.** Relative amounts of 18S and 25S RNA in the *rps24* mutants. Agilent 2100 Bioanalyzer electropherogram profiles of total RNA from Col-0, *rps24a-1*, *rps24b-2*, L*er* and *api6*. Total RNA was extracted from seedlings collected 15 das.

**Supplemental Figure 7.** Genetic interactions between *rps24b-2* and *rrp7-1*. Rosettes of Col-0 (A), *rps24b-2* (B), *rrp7-1* (C) and *rps24b-2 rrp7-1* plants (D). Photographs were taken from seedlings collected 14 das. Scale bars, 2 mm.

**Supplemental Figure 8.** Subcellular localization of 5.8S rRNA precursors. (A-A1) RNA-FISH assay in palisade mesophyll cells from first-node leaves of L*er* (A-C), *smo4-3* (D-F), *api6* (G-I), *rps24b-2 smo4-3* (J-L), Col-0 (M-O), *mtr4-2* (P-R), *rps24a-1* (S-U), *rps24b-2 mtr4-2* (V-X) and *rps24b-2* plants (Y-A1). Fluorescent signals correspond to DAPI (in blue; A, D, G, J, M, P S, V, and Y), which was used as a nuclear marker; the S9 probe labeled with Cy3 (in red; B, E, H, K, N, Q, T, W, and Z); and (C, F, I, L, O, R, U, X, and A1) their overlay. Plants were collected 21 das. Scale bars, 49 µm. (A2 and A3) Nucleus (A2) and nucleolus (A3) areas measured as DAPI and S9 probe fluorescence, respectively. Asterisks indicate significant differences from the wild type and parental lines (indicated by color) in a Student’s *t*-test (\**P* < 0.0001).

**Supplemental Figure 9.** Nucleolus organization in *rps24* root cells. (A-I) Visualization by immunolocalization of the fibrillarin nuclear marker in root cells of Col-0 (A, D, and G), *rps24a-1* (B, E, and H) and *rps24b-2* plants (C, F, and I). Fluorescent signals correspond to DAPI (A-C), the secondary antibody conjugated with TRITC for fibrillarin detection (D-E) and their overlay (G-I). Immunolocalization was performed in at least 5 seedlings per genotype, collected 5 das.

## REFERENCES

Ayash M, Abukhalaf M, Thieme D, Proksch C, Heilmann M, Schattat MH, Hoehenwarter W (2021) LC-MS based draft map of the *Arabidopsis thaliana* nuclear proteome and protein import in pattern triggered immunity. Front Plant Sci 12: 744103

Barakat A, Szick-Miranda K, Chang IF, Guyot R, Blanc G, Cooke R, Delseny M, Bailey-Serres J (2001) The organization of cytoplasmic ribosomal protein genes in the Arabidopsis genome. Plant Physiol 127: 398–415

Berná G, Robles P, Micol JL (1999) A mutational analysis of leaf morphogenesis in *Arabidopsis thaliana*. Genetics 152: 729–742

Bohnsack KE, Henras AK, Nielsen H, Bohnsack MT (2022) Making ends meet: a universal driver of large ribosomal subunit biogenesis. Trends Biochem Sci 48: 213–215

Burger F, Daugeron MC, Linder P (2000) Dbp10p, a putative RNA helicase from *Saccharomyces cerevisiae*, is required for ribosome biogenesis. Nucleic Acids Res 28: 2315–2323

Burgute BD, Peche VS, Steckelberg AL, Glockner G, Gassen B, Gehring NH, Noegel AA (2014) NKAP is a novel RS-related protein that interacts with RNA and RNA binding proteins. Nucleic Acids Res 42: 3177–3193

Carroll AJ, Heazlewood JL, Ito J, Millar AH (2008) Analysis of the *Arabidopsis* cytosolic ribosome proteome provides detailed insights into its components and their post-translational modification. Mol Cell Proteomics 7: 347–369

Carvalho CM, Santos AA, Pires SR, Rocha CS, Saraiva DI, Machado JP, Mattos EC, Fietto LG, Fontes EP (2008) Regulated nuclear trafficking of rpL10A mediated by NIK1 represents a defense strategy of plant cells against virus. PLOS Pathog 4: e1000247

Casanova-Sáez R, Candela H, Micol JL (2014) Combined haploinsufficiency and purifying selection drive retention of *RPL36a* paralogs in Arabidopsis. Sci Rep 4: 4122

Creff A, Sormani R, Desnos T (2010) The two Arabidopsis *RPS6* genes, encoding for cytoplasmic ribosomal proteins S6, are functionally equivalent. Plant Mol Biol 73: 533–546

Curtis MD, Grossniklaus U (2003) A Gateway cloning vector set for high-throughput functional analysis of genes in planta. Plant Physiol 133: 462–469

Chang IF, Szick-Miranda K, Pan S, Bailey-Serres J (2005) Proteomic characterization of evolutionarily conserved and variable proteins of Arabidopsis cytosolic ribosomes. Plant Physiol 137: 848–862

Chen HK, Pai CY, Huang JY, Yeh NH (1999) Human Nopp140, which interacts with RNA polymerase I: implications for rRNA gene transcription and nucleolar structural organization. Mol Cell Biol 19: 8536–8546

Choesmel V, Fribourg S, Aguissa-Toure AH, Pinaud N, Legrand P, Gazda HT, Gleizes PE (2008) Mutation of ribosomal protein RPS24 in Diamond-Blackfan anemia results in a ribosome biogenesis disorder. Hum Mol Genet 17: 1253–1263

Degenhardt RF, Bonham-Smith PC (2008) Arabidopsis ribosomal proteins RPL23aA and RPL23aB are differentially targeted to the nucleolus and are disparately required for normal development. Plant Physiol 147: 128–142

Durut N, Abou-Ellail M, Pontvianne F, Das S, Kojima H, Ukai S, de Bures A, Comella P, Nidelet S, Rialle S, Merret R, Echeverria M, Bouvet P, Nakamura K, Saez-Vasquez J (2014) A duplicated NUCLEOLIN gene with antagonistic activity is required for chromatin organization of silent 45S rDNA in Arabidopsis. Plant Cell 26: 1330–1344

Farooq M, Lindbaek L, Krogh N, Doganli C, Keller C, Monnich M, Goncalves AB, Sakthivel S, Mang Y, Fatima A, Andersen VS, Hussain MS, Eiberg H, Hansen L, Kjaer KW, Gopalakrishnan J, Pedersen LB, Mollgard K, Nielsen H, Baig SM, Tommerup N, Christensen ST, Larsen LA (2020) RRP7A links primary microcephaly to dysfunction of ribosome biogenesis, resorption of primary cilia, and neurogenesis. Nat Commun 11: 5816

Ferreira-Cerca S, Poll G, Gleizes PE, Tschochner H, Milkereit P (2005) Roles of eukaryotic ribosomal proteins in maturation and transport of pre-18S rRNA and ribosome function. Mol Cell 20: 263–275

Ferreyra ML, Biarc J, Burlingame AL, Casati P (2010a) Arabidopsis L10 ribosomal proteins in UV-B responses. Plant Signal Behav 5: 1222–1225

Ferreyra MLF, Pezza A, Biarc J, Burlingame AL, Casati P (2010b) Plant L10 ribosomal proteins have different roles during development and translation under ultraviolet-B stress. Plant Physiol 153: 1878–1894

Fujikura U, Horiguchi G, Ponce MR, Micol JL, Tsukaya H (2009) Coordination of cell proliferation and cell expansion mediated by ribosome-related processes in the leaves of *Arabidopsis thaliana*. Plant J 59: 499–508

Gazda HT, Grabowska A, Merida-Long LB, Latawiec E, Schneider HE, Lipton JM, Vlachos A, Atsidaftos E, Ball SE, Orfali KA, Niewiadomska E, Da Costa L, Tchernia G, Niemeyer C, Meerpohl JJ, Stahl J, Schratt G, Glader B, Backer K, Wong C, Nathan DG, Beggs AH, Sieff CA (2006) Ribosomal protein S24 gene is mutated in Diamond-Blackfan anemia. Am J Hum Genet 79: 1110–1118

Giavalisco P, Wilson D, Kreitler T, Lehrach H, Klose J, Gobom J, Fucini P (2005) High heterogeneity within the ribosomal proteins of the *Arabidopsis thaliana* 80S ribosome. Plant Mol Biol 57: 577–591

Hang R, Wang Z, Yang C, Luo L, Mo B, Chen X, Sun J, Liu C, Cao X (2021) Protein arginine methyltransferase 3 fine-tunes the assembly/disassembly of pre-ribosomes to repress nucleolar stress by interacting with RPS2B in *Arabidopsis*. Mol Plant 14: 223–236

Hawley RS, Gilliland WD (2006) Sometimes the result is not the answer: the truths and the lies that come from using the complementation test. Genetics 174: 5–15

Hoang T, Peng WT, Vanrobays E, Krogan N, Hiley S, Beyer AL, Osheim YN, Greenblatt J, Hughes TR, Lafontaine DL (2005) Esf2p, a U3-associated factor required for small-subunit processome assembly and compaction. Molecular and Cell Biology 25: 5523–5534

Horiguchi G, Fujikura U, Ferjani A, Ishikawa N, Tsukaya H (2006) Large-scale histological analysis of leaf mutants using two simple leaf observation methods: identification of novel genetic pathways governing the size and shape of leaves. Plant J 48: 638–644

Horiguchi G, Mollá-Morales A, Pérez-Pérez JM, Kojima K, Robles P, Ponce MR, Micol JL, Tsukaya H (2011) Differential contributions of ribosomal protein genes to *Arabidopsis thaliana* leaf development. Plant J 65: 724–736

Hruz T, Laule O, Szabo G, Wessendorp F, Bleuler S, Oertle L, Widmayer P, Gruissem W, Zimmermann P (2008) Genevestigator v3: a reference expression database for the meta-analysis of transcriptomes. Adv Bioinf 2008: 420747

Hummel M, Dobrenel T, Cordewener JJ, Davanture M, Meyer C, Smeekens SJ, Bailey-Serres J, America TA, Hanson J (2015) Proteomic LC-MS analysis of Arabidopsis cytosolic ribosomes: Identification of ribosomal protein paralogs and re-annotation of the ribosomal protein genes. J Proteom 128: 436–449

Jiang Y, Wang D, Yao Y, Eubel H, Kunzler P, Moller IM, Xu D (2021) MULocDeep: a deep-learning framework for protein subcellular and suborganellar localization prediction with residue-level interpretation. Comput Struct Biotechnol J 19: 4825–4839

Kang J, Brajanovski N, Chan KT, Xuan J, Pearson RB, Sanij E (2021) Ribosomal proteins and human diseases: molecular mechanisms and targeted therapy. Signal Transduct Target Ther 6: 323

Khan A, Garbelli A, Grossi S, Florentin A, Batelli G, Acuna T, Zolla G, Kaye Y, Paul LK, Zhu JK, Maga G, Grafi G, Barak S (2014) The Arabidopsis STRESS RESPONSE SUPPRESSOR DEAD-box RNA helicases are nucleolar- and chromocenter-localized proteins that undergo stress-mediated relocalization and are involved in epigenetic gene silencing. Plant J 79: 28–43

Kim YK, Kim S, Shin YJ, Hur YS, Kim WY, Lee MS, Cheon CI, Verma DP (2014) Ribosomal protein S6, a target of rapamycin, is involved in the regulation of rRNA genes by possible epigenetic changes in *Arabidopsis*. J Biol Chem 289: 3901–3912

Kojima H, Suzuki T, Kato T, Enomoto K, Sato S, Kato T, Tabata S, Sáez-Vásquez J, Echeverría M, Nakagawa T, Ishiguro S, Nakamura K (2007) Sugar-inducible expression of the nucleolin-1 gene of *Arabidopsis thaliana* and its role in ribosome synthesis, growth and development. Plant J 49: 1053–1063

Lange H, Sement FM, Gagliardi D (2011) MTR4, a putative RNA helicase and exosome co-factor, is required for proper rRNA biogenesis and development in *Arabidopsis thaliana*. Plant J 68: 51–63

Li S, Liu K, Zhang S, Wang X, Rogers K, Ren G, Zhang C, Yu B (2017) STV1, a ribosomal protein, binds primary microRNA transcripts to promote their interaction with the processing complex in *Arabidopsis*. Proc Natl Acad Sci USA 114: 1424–1429

Martinez-Seidel F, Beine-Golovchuk O, Hsieh YC, Kopka J (2020) Systematic review of plant ribosome heterogeneity and specialization. Front Plant Sci 11: 948

Meinke DW (2013) A survey of dominant mutations in *Arabidopsis thaliana*. Trends Plant Sci 18: 84–91

Mergner J, Frejno M, List M, Papacek M, Chen X, Chaudhary A, Samaras P, Richter S, Shikata H, Messerer M, Lang D, Altmann S, Cyprys P, Zolg DP, Mathieson T, Bantscheff M, Hazarika RR, Schmidt T, Dawid C, Dunkel A, Hofmann T, Sprunck S, Falter-Braun P, Johannes F, Mayer KFX, Jurgens G, Wilhelm M, Baumbach J, Grill E, Schneitz K, Schwechheimer C, Kuster B (2020) Mass-spectrometry-based draft of the Arabidopsis proteome. Nature 579: 409–414

Micol-Ponce R, Sarmiento-Mañús R, Fontcuberta-Cervera S, Cabezas-Fuster A, de Bures A, Sáez-Vásquez J, Ponce MR (2020) SMALL ORGAN4 is a ribosome biogenesis factor involved in 5.8S ribosomal RNA maturation. Plant Physiol 184: 2022–2039

Micol-Ponce R, Sarmiento-Mañús R, Ruiz-Bayón A, Montacié C, Sáez-Vásquez J, Ponce MR (2018) Arabidopsis RIBOSOMAL RNA PROCESSING7 is required for 18S rRNA maturation. Plant Cell 30: 2855–2872

Montacié C, Durut N, Opsomer A, Palm D, Comella P, Picart C, Carpentier MC, Pontvianne F, Carapito C, Schleiff E, Sáez-Vásquez J (2017) Nucleolar proteome analysis and proteasomal activity assays reveal a link between nucleolus and 26S proteasome in *A. thaliana*. Front Plant Sci 8: 1815

Navarro-Quiles C, Mateo-Bonmatí E, Candela H, Robles P, Martínez-Laborda A, Fernández Y, Simura J, Ljung K, Rubio V, Ponce MR, Micol JL (2022) The Arabidopsis ATP-Binding Cassette E protein ABCE2 is a conserved component of the translation machinery. Front Plant Sci 13: 1009895

Nguyen TH, Li J, Galej WP, Oshikane H, Newman AJ, Nagai K (2013) Structural basis of Brr2-Prp8 interactions and implications for U5 snRNP biogenesis and the spliceosome active site. Structure 21: 910–919

Ni C, Buszczak M (2023) The homeostatic regulation of ribosome biogenesis. Semin Cell Dev Biol 136: 13–26

Obayashi T, Hibara H, Kagaya Y, Aoki Y, Kinoshita K (2022) ATTED-II v11: A plant gene coexpression database using a sample balancing technique by subagging of principal components. Plant Cell Physiol 63: 869–881

Pajerowski AG, Nguyen C, Aghajanian H, Shapiro MJ, Shapiro VS (2009) NKAP is a transcriptional repressor of notch signaling and is required for T cell development. Immunity 30: 696–707

Palm D, Simm S, Darm K, Weis BL, Ruprecht M, Schleiff E, Scharf C (2016) Proteome distribution between nucleoplasm and nucleolus and its relation to ribosome biogenesis in *Arabidopsis thaliana*. RNA Biol 13: 441–454

Palm D, Streit D, Shanmugam T, Weis BL, Ruprecht M, Simm S, Schleiff E (2019) Plant-specific ribosome biogenesis factors in Arabidopsis thaliana with essential function in rRNA processing. Nucleic Acids Res 47: 1880–1895

Pan WA, Tsai HY, Wang SC, Hsiao M, Wu PY, Tsai MD (2015) The RNA recognition motif of NIFK is required for rRNA maturation during cell cycle progression. RNA Biol 12: 255–267

Parry G, Ward S, Cernac A, Dharmasiri S, Estelle M (2006) The Arabidopsis SUPPRESSOR OF AUXIN RESISTANCE proteins are nucleoporins with an important role in hormone signaling and development. Plant Cell 18: 1590–1603

Pasternak T, Tietz O, Rapp K, Begheldo M, Nitschke R, Ruperti B, Palme K (2015) Protocol: an improved and universal procedure for whole-mount immunolocalization in plants. Plant Methods 11: 50

Pendle AF, Clark GP, Boon R, Lewandowska D, Lam YW, Andersen J, Mann M, Lamond AI, Brown JW, Shaw PJ (2005) Proteomic analysis of the Arabidopsis nucleolus suggests novel nucleolar functions. Mol Biol Cell 16: 260–269

Pérez-Pérez JM, Candela H, Micol JL (2009) Understanding synergy in genetic interactions. Trends Genet 25: 368–376

Petibon C, Malik Ghulam M, Catala M, Abou Elela S (2021) Regulation of ribosomal protein genes: an ordered anarchy. WIREs RNA 12: e1632

Petricka JJ, Nelson TM (2007) Arabidopsis nucleolin affects plant development and patterning. Plant Physiol 144: 173–186

Ponce MR, Robles P, Lozano FM, Brotóns MA, Micol JL (2006) Low-resolution mapping of untagged mutations. Methods Mol Biol 323: 105–113

Ponce MR, Robles P, Micol JL (1999) High-throughput genetic mapping in *Arabidopsis thaliana*. Mol Gen Genet 261: 408–415

Pontvianne F, Abou-Ellail M, Douet J, Comella P, Matia I, Chandrasekhara C, Debures A, Blevins T, Cooke R, Medina FJ, Tourmente S, Pikaard CS, Sáez-Vásquez J (2010) Nucleolin is required for DNA methylation state and the expression of *rRNA* gene variants in *Arabidopsis thaliana*. PLOS Genet 6: e1001225

Pontvianne F, Matia I, Douet J, Tourmente S, Medina FJ, Echeverria M, Sáez-Vásquez J (2007) Characterization of *AtNUC-L1* reveals a central role of nucleolin in nucleolus organization and silencing of *AtNUC-L2* gene in Arabidopsis. Mol Biol Cell 18: 369–379

Robledo S, Idol RA, Crimmins DL, Ladenson JH, Mason PJ, Bessler M (2008) The role of human ribosomal proteins in the maturation of rRNA and ribosome production. RNA 14: 1918–1929

Sáez-Vásquez J, Delseny M (2019) Ribosome biogenesis in plants: from functional 45S ribosomal DNA organization to ribosome assembly factors. Plant Cell 31: 1945–1967

Salih KJ, Duncan O, Li L, Trosch J, Millar AH (2020) The composition and turnover of the *Arabidopsis thaliana* 80S cytosolic ribosome. Biochem J 477: 3019–3032

Samaha H, Delorme V, Pontvianne F, Cooke R, Delalande F, Van Dorsselaer A, Echeverria M, Saez-Vasquez J (2010) Identification of protein factors and U3 snoRNAs from a Brassica oleracea RNP complex involved in the processing of pre-rRNA. Plant J 61: 383–398

Sánchez-García AB, Aguilera V, Micol-Ponce R, Jover-Gil S, Ponce MR (2015) Arabidopsis *MAS2*, an essential gene that encodes a homolog of animal NF-κB activating protein, is involved in 45S ribosomal DNA silencing. Plant Cell 27: 1999–2015

Savada RP, Bonham-Smith PC (2014) Differential transcript accumulation and subcellular localization of Arabidopsis ribosomal proteins. Plant Sci 223: 134–145

Scarpin MR, Busche M, Martinez RE, Harper LC, Reiser L, Szakonyi D, Merchante C, Lan T, Xiong W, Mo B, Tang G, Chen X, Bailey-Serres J, Browning KS, Brunkard JO (2022) An updated nomenclature for plant ribosomal protein genes. Plant Cell 35: 640–643

Scott MS, Troshin PV, Barton GJ (2011) NoD: a nucleolar localization sequence detector for eukaryotic and viral proteins. BMC Bioinformatics 12: 317

Schwacke R, Schneider A, van der Graaff E, Fischer K, Catoni E, Desimone M, Frommer WB, Flugge UI, Kunze R (2003) ARAMEMNON, a novel database for Arabidopsis integral membrane proteins. Plant Physiol 131: 16–26

Shanmugam T, Streit D, Schroll F, Kovacevic J, Schleiff E (2021) Dynamics and thermal sensitivity of ribosomal RNA maturation paths in plants. J Exp Bot 72: 7626–7644

Slavov N, Semrau S, Airoldi E, Budnik B, van Oudenaarden A (2015) Differential stoichiometry among core ribosomal proteins. Cell Rep 13: 865–873

Sperschneider J, Catanzariti AM, DeBoer K, Petre B, Gardiner DM, Singh KB, Dodds PN, Taylor JM (2017) LOCALIZER: subcellular localization prediction of both plant and effector proteins in the plant cell. Sci Rep 7: 44598

Tafforeau L, Zorbas C, Langhendries JL, Mullineux ST, Stamatopoulou V, Mullier R, Wacheul L, Lafontaine DL (2013) The complexity of human ribosome biogenesis revealed by systematic nucleolar screening of pre-rRNA processing factors. Mol Cell 51: 539–551

Thoms M, Thomson E, Bassler J, Gnadig M, Griesel S, Hurt E (2015) The exosome is recruited to RNA substrates through specific adaptor proteins. Cell 162: 1029–1038

Tyagi A, Gupta A, Dutta A, Potluri P, Batti B (2020) A review of Diamond-Blackfan anemia: current evidence on involved genes and treatment modalities. Cureus 12: e10019

Wang R, Zhao J, Jia M, Xu N, Liang S, Shao J, Qi Y, Liu X, An L, Yu F (2018) Balance between cytosolic and chloroplast translation affects leaf variegation. Plant Physiol 176: 804–818

Warner JR (1999) The economics of ribosome biosynthesis in yeast. Trends Biochem Sci 24: 437–440

Weaver PL, Sun C, Chang TH (1997) Dbp3p, a putative RNA helicase in *Saccharomyces cerevisiae*, is required for efficient pre-rRNA processing predominantly at site A_3_. Mol Cell Biol 17: 1354–1365

Wilson DN, Cate JHD (2012) The structure and function of the eukaryotic ribosome. Cold Spring Harb Perspect Biol 4: a011536

Xiong W, Zhang J, Lan T, Kong W, Wang X, Liu L, Chen X, Mo B (2021) High resolution RNA-seq profiling of genes encoding ribosomal proteins across different organs and developmental stages in *Arabidopsis thaliana*. Plant Direct 5: e00320

Yang Y, Isaac C, Wang C, Dragon F, Pogacic V, Meier UT (2000) Conserved composition of mammalian box H/ACA and box C/D small nucleolar ribonucleoprotein particles and their interaction with the common factor Nopp140. Mol Biol Cell 11: 567–577

Zhu P, Wang Y, Qin N, Wang F, Wang J, Deng XW, Zhu D (2016) Arabidopsis small nucleolar RNA monitors the efficient pre-rRNA processing during ribosome biogenesis. Proc Natl Acad Sci U S A 113: 11967–11972

