## Supplemental Figures and Tables for "Cross-kingdom conservation of Arabidopsis RPS24 function in 18S rRNA maturation"

### Supporting Figures, Tables and References

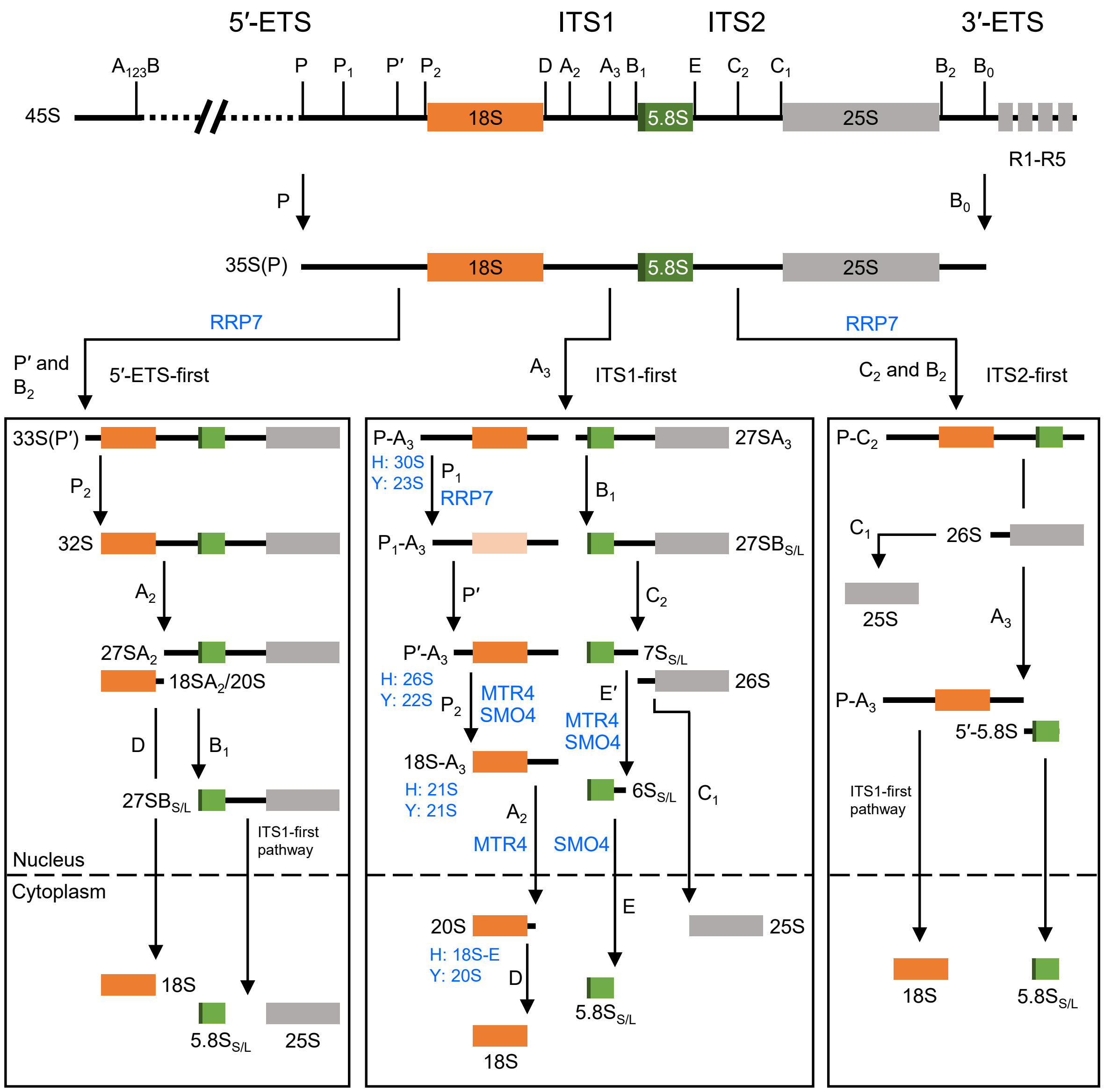


**Supplemental Figure 1.** Overview of 45S pre-rRNA processing in Arabidopsis. Colored boxes represent the sequences of the three mature rRNAs transcribed from the 45S rDNA genes. Vertical arrows indicate endonucleolytic cleavages; letters indicate the cleavage site in the corresponding pre-rRNA. Only the relevant factors and the human (H) and yeast (Y) 18S pre-rRNAs relevant to this study are represented. Based on information from Sáez-Vasquez and Delseny (2019).

```
RPS24B         1  MAEKAVTIRTRNFMTNRLLARKQFVIDVLHPGRANVSKAELKEKLARMYEVKDPNAIFCF
RPS24A         1  MAEKAVTIRTRKFMTNRLLSRKQFVIDVLHPGRANVSKAELKEKLARMYEVKDPNAIFVF
consensus      1  *********** ******* ************************************** *

RPS24B        61  KFRTHFGGGKSSGYGLIYDTVENAKKFEPKYRLIRNGLDTKIEKSRKQIKERKNRAKKIR
RPS24A        61  KFRTHFGGGKSSGFGLIYDTVESAKKFEPKYRLIRNGLDTKIEKSRKQIKERKNRAKKIR
consensus     61  *************.******** *************************************

RPS24B       121  GVKKTKAGDTKKK
RPS24A       121  GVKKTKAGDAKKK
consensus    121  ********* ***
```

**Supplemental Figure 2.** Sequence conservation between the RPS24A and RPS24B paralogs. Amino acid sequence alignment of Arabidopsis RPS24B and RPS24A. The 14 amino acids that are absent from the RPS24B variant produced by the expression of the *api6* allele are highlighted in red letters. Identical and similar residues are shaded in black and gray, respectively. Asterisks and dots in the consensus line indicate identical and conserved residues, respectively. Numbers indicate residue positions. The alignment was obtained using ClustalW2 and shaded with Boxshade 3.21 (http://www.ch.embnet.org/software/BOX_form.html). Nuclear and nucleolar localization sequences predicted with LOCALIZER (https://localizer.csiro.au/) and NoD software (http://www.compbio.dundee.ac.uk/www-nod/index.jsp) are underlined in black and green, respectively.

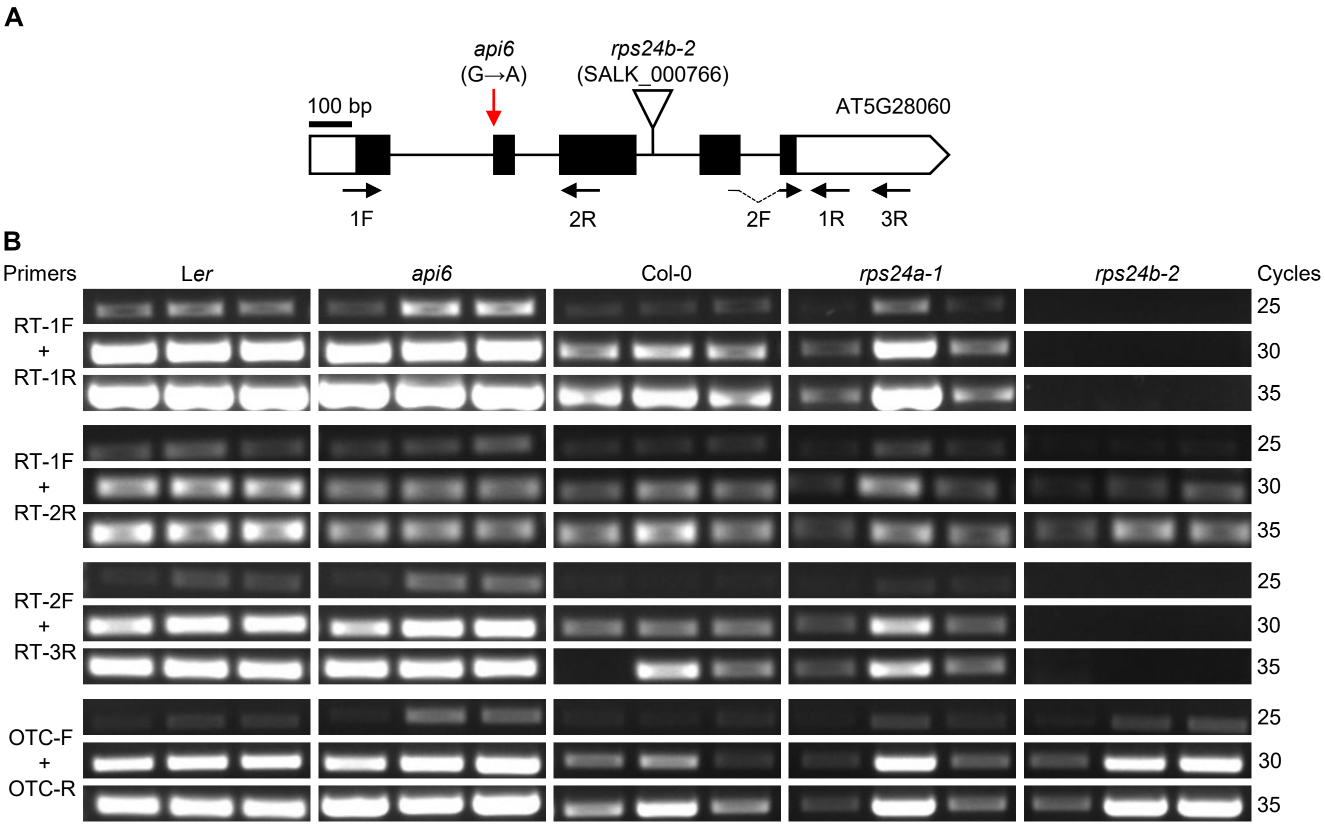


**Supplemental Figure 3.** *RPS24B* expression analysis. (A) Schematic representation of *RPS24B* and its mutant alleles used in this study. Gene structure is represented as described in the legend of Figure 1. Black arrows represent the primers used for semiquantitative RT-PCR amplifications in (B). (B) Semiquantitative RT-PCR analysis of *RPS24B* in the *rps24a* and *rps24b* mutants used in this study. The bands for each PCR were visualized after 25, 30 and 35 cycles of amplification. Total RNA was extracted from seedlings collected at 15 das. Transcripts from the *OTC* gene were used as an internal control. The primer sequences used are shown in Supplemental Table 2.

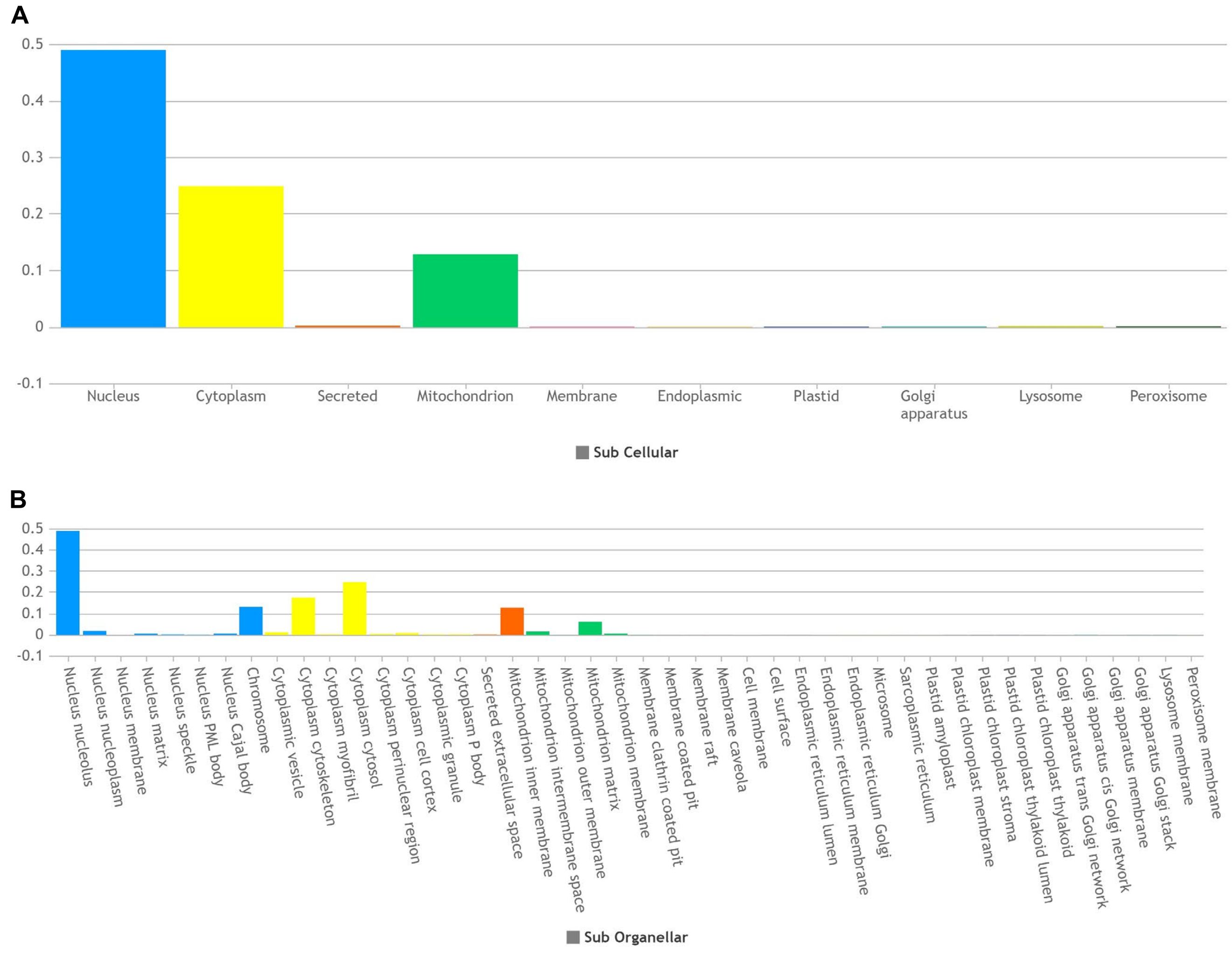


**Supplemental Figure 4.** Predicted localization of RPS24A. (A-B) Predicted subcellular (A) and sub-organellar (B) localization of RPS24A using MULocDeep software (https://mu-loc.org/).

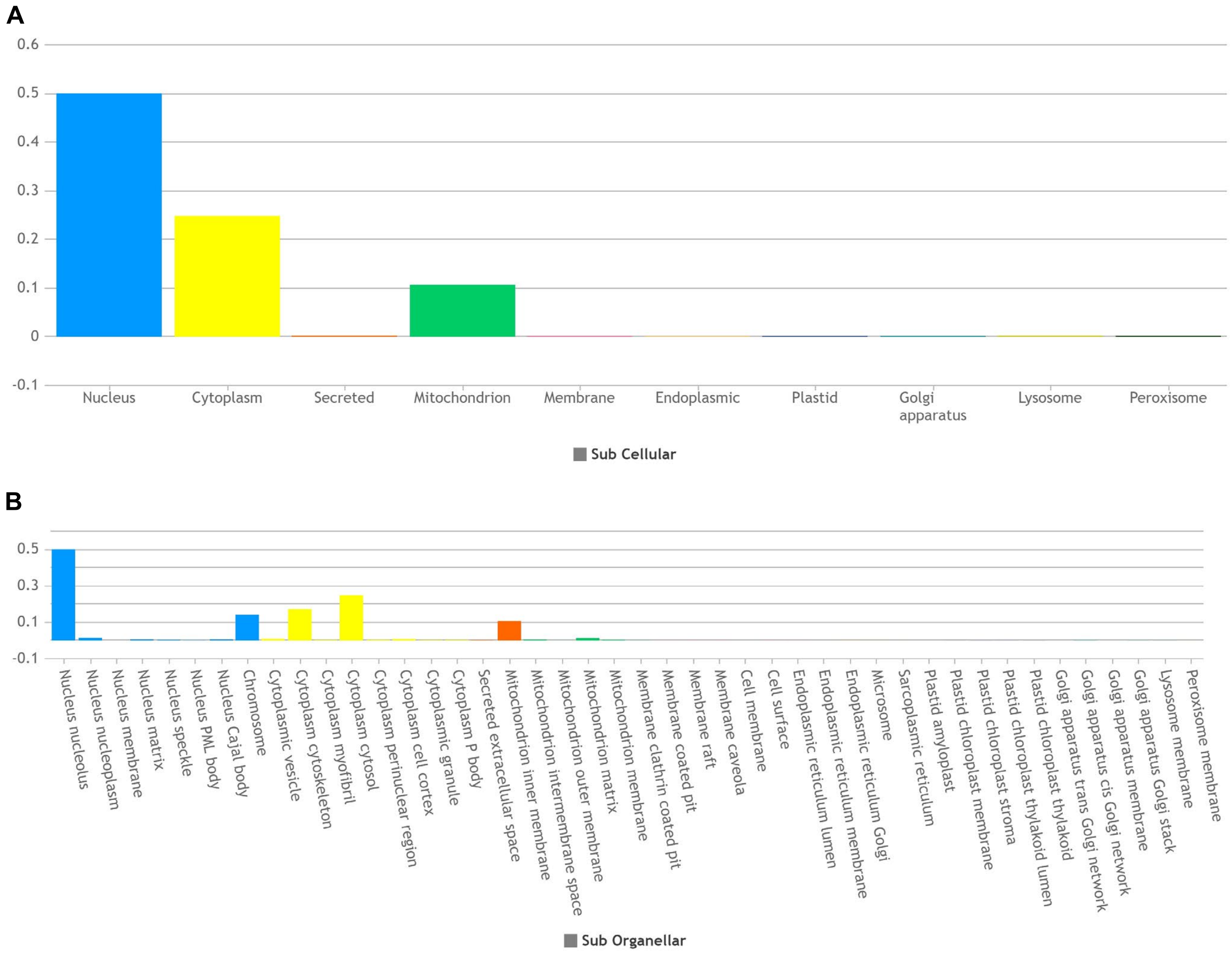


**Supplemental Figure 5.** Predicted localization of RPS24B. (A-B) Predicted subcellular (A) and sub-organellar (B) localization of RPS24B protein using MULocDeep software (https://mu-loc.org/).

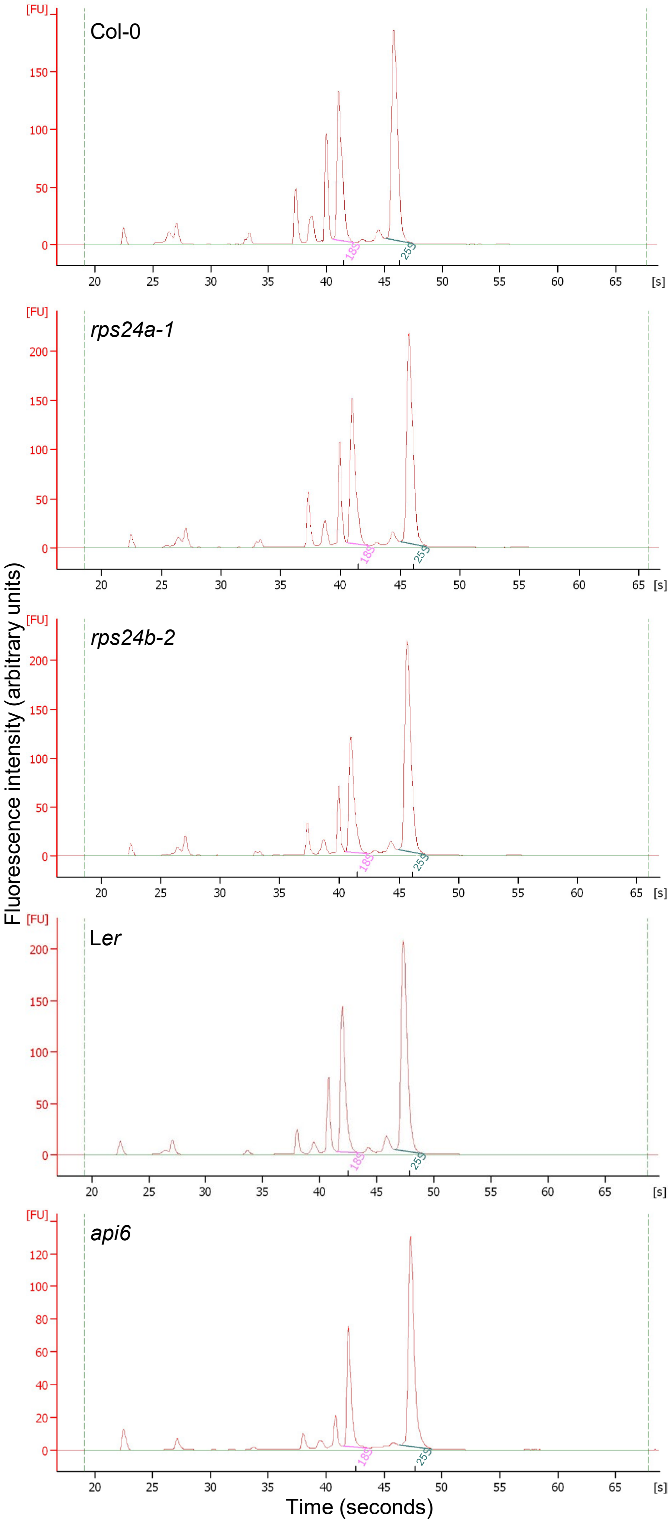


**Supplemental Figure 6.** Relative amounts of 18S and 25S RNA in the *rps24* mutants. Agilent 2100 Bioanalyzer electropherogram profiles of total RNA from Col-0, *rps24a-1*, *rps24b-2*, L*er* and *api6*. Total RNA was extracted from seedlings collected 15 das.

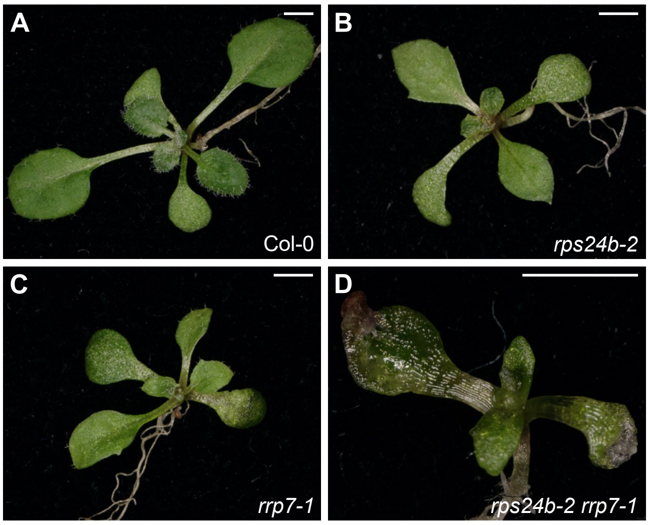


**Supplemental Figure 7.** Genetic interactions between *rps24b-2* and *rrp7-1*. Rosettes of Col-0 (A), *rps24b-2* (B), *rrp7-1* (C) and *rps24b-2 rrp7-1* plants (D). Photographs were taken from seedlings collected 14 das. Scale bars, 2 mm.

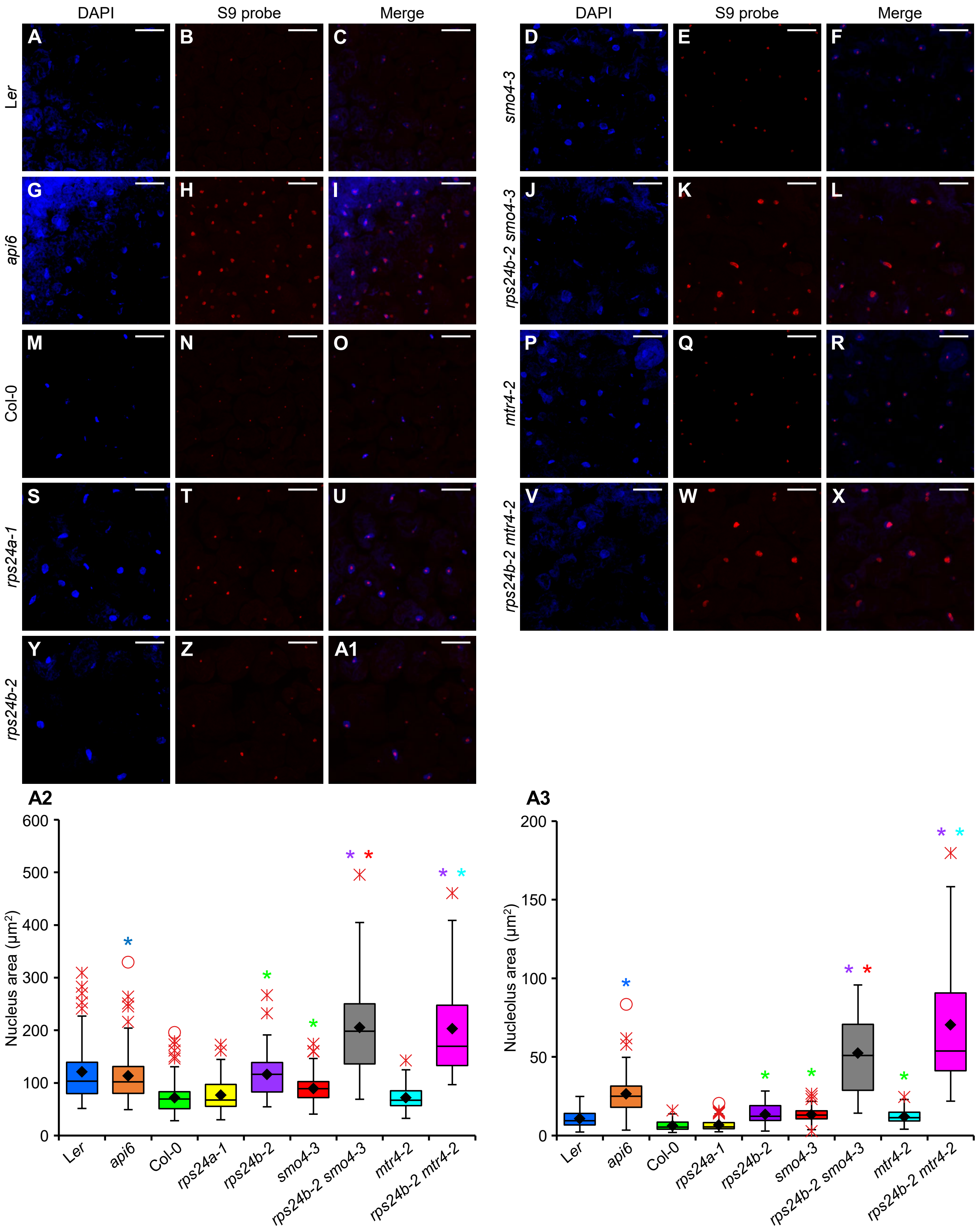

DAPI
S9 probe
Merge
A
B
C
Ler
D
E
F
smo4-3
G
H
I
api6
J
K
L
rps24b-2 smo4-3
M
N
O
Col-0
P
Q
R
mtr4-2
S
T
U
rps24a-1
V
W
X
rps24b-2 mtr4-2
Y
Z
A1
rps24b-2
A2
Nucleus area (μm²)
0
100
200
300
400
500
600
Ler
api6
Col-0
rps24a-1
rps24b-2
smo4-3
rps24b-2 smo4-3
mtr4-2
rps24b-2 mtr4-2
A3
Nucleolus area (μm²)
0
50
100
150
200
Ler
api6
Col-0
rps24a-1
rps24b-2
smo4-3
rps24b-2 smo4-3
mtr4-2
rps24b-2 mtr4-2

**Supplemental Figure 8.** Subcellular localization of 5.8S rRNA precursors. (A-A1) RNA-FISH assay in palisade mesophyll cells from first-node leaves of L*er* (A-C), *smo4-3* (D-F), *api6* (G-I), *rps24b-2 smo4-3* (J-L), Col-0 (M-O), *mtr4-2* (P-R), *rps24a-1* (S-U), *rps24b-2 mtr4-2* (V-X) and *rps24b-2* plants (Y-A1). Fluorescent signals correspond to DAPI (in blue; A, D, G, J, M, P S, V, and Y), which was used as a nuclear marker; the S9 probe labeled with Cy3 (in red; B, E, H, K, N, Q, T, W, and Z); and (C, F, I, L, O, R, U, X, and A1) their overlay. Plants were collected 21 das. Scale bars, 49 µm. (A2 and A3) Nucleus (A2) and nucleolus (A3) areas measured as DAPI and S9 probe fluorescence, respectively. Asterisks indicate significant differences from the wild type and parental lines (indicated by color) in a Student's *t*-test (**P* < 0.0001).

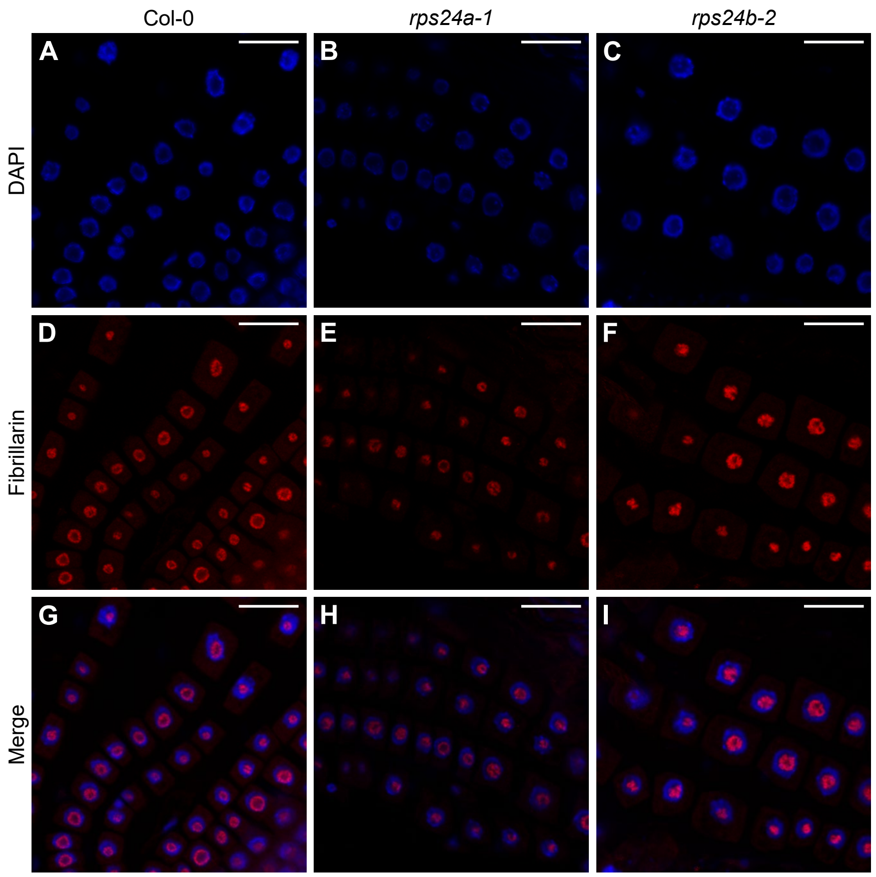


**Supplemental Figure 9.** Nucleolus organization in *rps24* root cells. (A-I) Visualization by immunolocalization of the fibrillarin nuclear marker in root cells of Col-0 (A, D, and G), *rps24a-1* (B, E, and H) and *rps24b-2* plants (C, F, and I). Fluorescent signals correspond to DAPI (A-C), the secondary antibody conjugated with TRITC for fibrillarin detection (D-E) and their overlay (G-I). Immunolocalization was performed in at least 5 seedlings per genotype, collected 5 das.

**Supplemental Table 1.** Primer sets used for the fine mapping of *api6*

| Marker name | Locus | Oligonucleotide sequences (5′→3′) | | PCR product size (bp) | |
|---|---|---|---|---|---|
| | | Forward primer | Reverse primer | L*er* | Col-0 |
| cer449133 | AT5G27905-AT5G27910 | CGATTGTTTGTTTCAACTTTCAA[a] | GCGTATGTTTGGGGGATAGG[b] | 262 | 287 |
| cer451402 | AT5G28200 | GTGTATCATGGACCATCGCG[c] | AATTGTTTTGTAGGTCGCTAAC[d] | 237 | 229 |

The oligonucleotide names are: [a]cer449133-F, [b]cer449133-R, [c]cer451402-F and [d]cer451402-R.

**Supplemental Table 2.** Other primers used in this work

| Purpose | | Oligonucleotide name(s) | Oligonucleotide sequences (5→3′) | |
|---|---|---|---|---|
| | | | Forward primer (F) | Reverse primer (R) |
| Genotyping of | *rps24b-2*/*api6* | AT5G28060-F1/R1 | CTAATCTATTCTCTGGGCATGG | CAGCCTTGGTCTACACTCAC |
| | *rps24a-1* | AT3G04920-F1/R1 | CCTGGAAGAGCCAATGTTTCA | ATGGGAATGGTGGAAAGAGAC |
| | *smo4-3* | NOP53-F1/R1[a] | GTCTCGAACTTTTTCCTTGGG | AGTATTCCTCGCTTCTCGAGG |
| | *mtr4-2* | MTR4-F/R[a] | TTTGTCAATACCTCGACGTCC | ATTGTCTGCGTACTGTGGGTC |
| | *rrp7-1* | RRP7-F1/R1 | CTCATGAAGAACGCCTTGAAC | GTGGAGATCGTGGAGATGAAG |
| | *parl1-2* | PARL1-F/R[b] | AGTTGCTGTCACCAAGAAG | TGGCCTACCATGGAATTCA |
| T-DNA insertion verification | | Salk_LBb1.3[c] | GCGTGGACCGCTTGCTGCAACT | |
| | | Salk_Rb1[c] | CGTGACTCCCTTAATTCTCCGC | |
| | | Sail_LB1[c] | GCCTTTTCAGAAATGGATAAATAGCCTTGCTTCC | |
| Construction of transgenes | | 35Spro:RPS24B:GFP-F/R | *GGGGACCACTTTGTACAAGAAAGCTGGGTG*TTACTTCTTCTTGGTATCACCAGC | *GGGGACCACTTTGTACAAGAAAGCTGGGTG*CTTCTTCTTGGTATCACCAGC |
| Semiquantitative RT-PCR | | RPS24B-RT-1F/1R | AAGTAAATCGCAGCCATGGC | TTGTTCTGCATCACTCCTTCTT |
| | | RPS24B-RT-2F/2R | AGTACAGACTTATCAGGAATGGA | GGCGAGGATGTATGAGGTTAAG |
| | | RPS24B-RT-3R | | CCTCTTGCGTTTCGGAGATT |
| | | OTC-REV/3D | GCATGCATGCGATTCTCCGC | TCCTTGCCAAATCATGGCCG |
| Quantitative RT-PCR | | 45S pre-rRNA-45S-F/R[d] | CGGTCGGTCATTCCTCGTGTCGATATC | TATAGGGGGGTGGGTGTTGAGGGA |
| 45S rDNA variant expression | | p3/p4[e] | GACAGACTTGTCCAAAACGCCCACC | CTGGTCGAGGAATCCTGGACGATT |
| | | OTC-F/R[f] | TGAAGGGACAAAGGTTGTGTATGTT | CGCAGACAAGTGGAATGGA |
| Probes for RNA gel blots and *in situ* hybridization | | S2-F/R[g] | TAGGCTGTCCCGAAGTATC[h] | TCACTTCGAGTCACCGTCGACA[j] |
| | | S7[g] | GTCGTTCTGTTTTGGACAGGTATCGA[h] | |
| | | S9[g] | AGGATGGTGAGGGACGACGATTT[h,i] | |

Sequences taken from [a]Micol-Ponce et al., 2020, [b]Micol-Ponce et al., 2018, [c]http://signal.salk.edu/tdnaprimers.2.html, [d]Zhu et al., 2016, [e]Pontvianne et al., 2010, [f]Cnops et al., 2004, and [g]Lange et al., 2011. [h,i]Oligonucleotides labeled with [h]DIG (Digoxigenin) and [i]Cy3 (Cyanine 3). The *att*B sequences are shown in italics.

### SUPPLEMENTAL REFERENCES


**Cnops G, Jover-Gil S, Peters JL, Neyt P, De Block S, Robles P, Ponce MR, Gerats T, Micol JL, Van Lijsebettens M** (2004) The rotunda2 mutants identify a role for the *LEUNIG* gene in vegetative leaf morphogenesis. J Exp Bot **55:** 1529-1539

**Lange H, Sement FM, Gagliardi D** (2011) MTR4, a putative RNA helicase and exosome co-factor, is required for proper rRNA biogenesis and development in *Arabidopsis thaliana*. Plant J **68:** 51-63

**Micol-Ponce R, Sarmiento-Mañús R, Fontcuberta-Cervera S, Cabezas-Fuster A, de Bures A, Sáez-Vásquez J, Ponce MR** (2020) SMALL ORGAN4 is a ribosome biogenesis factor involved in 5.8S ribosomal RNA maturation. Plant Physiol **184:** 2022-2039

**Micol-Ponce R, Sarmiento-Mañús R, Ruiz-Bayón A, Montacié C, Sáez-Vásquez J, Ponce MR** (2018) Arabidopsis RIBOSOMAL RNA PROCESSING7 is required for 18S rRNA maturation. The Plant Cell **30:** 2855-2872

**Pontvianne F, Abou-Ellail M, Douet J, Comella P, Matia I, Chandrasekhara C, Debures A, Blevins T, Cooke R, Medina FJ, Tourmente S, Pikaard CS, Sáez-Vásquez J** (2010) Nucleolin is required for DNA methylation state and the expression of *rRNA* gene variants in *Arabidopsis thaliana*. PLOS Genet **6:** e1001225

**Zhu P, Wang Y, Qin N, Wang F, Wang J, Deng XW, Zhu D** (2016) Arabidopsis small nucleolar RNA monitors the efficient pre-rRNA processing during ribosome biogenesis. Proc Natl Acad Sci U S A **113:** 11967-11972
